# Bacterial metabolic signatures in MASLD predicted through gene-centric studies in stool metagenomes

**DOI:** 10.1101/2025.05.03.652020

**Authors:** Juan Manuel Medina-Méndez, Paula Iruzubieta, Raúl Fernández-López, Javier Crespo, Fernando de la Cruz

**Affiliations:** Instituto de Biomedicina y Biotecnología de Cantabria (IBBTEC), Universidad de Cantabria-CSIC, Santander, Spain; Servicio de Gastroenterología y Hepatología. Grupo de Investigación Clínica y Traslacional en Enfermedades Digestivas. Instituto de Investigación Valdecilla (IDIVAL). Hospital Universitario Marqués de Valdecilla, Santander, Spain

**Keywords:** MASLD, gut microbiome, metagenomic sample, metabolic gene, RPKSM, differential gene abundance

## Abstract

Metabolic dysfunction-associated steatotic liver disease (MASLD) is a multifactorial condition influenced by the gut microbiome (GM). While previous studies have reported inconsistent associations between MASLD and key microbial clades using low-resolution 16S rRNA profiling, we employed high-resolution metagenomic sequencing and multi-marker taxonomic classification across three independent cohorts to identify robust microbial and functional signatures of MASLD. We consistently detected a depletion of *Agathobacter rectalis*, a known butyrate producer, in MASLD patients. Functionally, MASLD was characterized by a depletion of genes involved in butyrate and methane biosynthesis-particularly within the crotonyl-butyryl-CoA axis-alongside an enrichment of genes driving the production of endogenous alcohols such as ethanol and 1-propanol. Genes encoding these fermentative pathways, often organized in operons like *pdu* and *tor*, were more abundant in MASLD samples, indicating a potential shift toward alcohol-producing metabolism. These geno-metabolic changes were accompanied by a broader displacement of beneficial taxa and an increase in accessory gene content across the GM, underscoring the limitations of taxonomy-based disease associations. Many of the differentially abundant genes were also found on plasmids, suggesting that horizontal gene transfer contributes to strain-level metabolic variability relevant to MASLD progression. Our findings support a model in which GM-driven metabolic shifts-rather than taxonomic changes alone-play a central role in MASLD pathogenesis, highlighting the importance of functional and mobile genetic element (MGEs) profiling for uncovering mechanistic links between the microbiome and liver disease.

## 1. Introduction

Metabolic dysfunction-associated steatotic liver disease (MASLD) is the most common liver disease worldwide, affecting approximately one quarter of the global population^1^. Clinically, MASLD is defined by the presence of steatosis (i.e., accumulation of intrahepatic triglycerides) in more than 5% of hepatocytes, in association with metabolic risk factors (particularly, obesity and type 2 diabetes) and in the absence of excessive alcohol consumption (≥30 g per day for men and ≥20 g per day for women) or other chronic liver diseases^2^. However, the term encompasses a spectrum of pathological situations, ranging from simple steatosis to metabolic dysfunction-associated steatohepatitis (MASH), which can further progress to fibrosis, cirrhosis and hepatocellular carcinoma^3^. MASLD also significantly increases the risk of cardiovascular disease, extrahepatic malignancies (e.g., colorectal or breast cancer), liver-related complications or chronic kidney disease^4^. The pathogenesis of MASLD is multifactorial, involving genetic, environmental, inflammatory and metabolic factors, and it has a bidirectional association with components of the metabolic syndrome, particularly insulin resistance^5^.

The gut microbiome (GM) interacts with the liver via the gut-liver axis (GLA) through the portal vein, which transports GM-derived metabolites to the liver. This axis can be disrupted by multiple interconnected factors, affecting the progression of MASLD and fibrosis^6^. Alterations of the GM composition and/or their associated metabolic production can lead to an increased production of harmful metabolites such as pathogen-associated molecular patterns, which induce inflammatory responses in the liver when they reach it through the bloodstream, leading to liver injury^7,8^. The transient of these endotoxins into portal circulation is facilitated by an increased intestinal permeability caused by a disruption of tight junction proteins in the intestinal membrane^9^. Additionally, alterations of the bile acid composition and circulation affect liver metabolism and function. Since bile acids play a role in the regulation of glucose and lipid metabolism, their dysregulation contributes to gut inflammation and MASLD^10,11^.

Several studies have attempted to establish a causal link between bacterial metabolism and MASLD^12,13^. However, metabolic levels can also be influenced by host metabolism, diet or environmental factors. Similarly, metataxonomic analysis have explored the relationship between GM composition and MASLD^14,15^. Although 16S rRNA gene sequencing provides taxonomic resolution at the genus or species level, it fails to capture strain-level metabolic variability. Bacterial genomic plasticity allows key metabolic genes, such as those potentially relevant to MASLD pathogenesis, to be encoded in their accessory genome, including MGEs, further complicating the association between taxonomy and metabolic function^16^.

In this study, we used the genes encoding metabolic enzymes involved in the final steps of microbial pathways leading to the production of butyrate, short-chain alcohols (SCAs) ethanol and propanol, methane, trimethylamine (TMA) and trimethylamine N-oxide (TMAO) as proxies to determine their abundance in MASLD. To achieve this, we isolated these gene families and quantified them in over 550 metagenomes obtained from fecal samples of MASLD and healthy patients, stratified across three cohorts. We also explored taxonomic signatures to gain a comprehensive view of bacterial diversity and abundance. Finally, we assessed the presence of these genes within the human GM genomes and plasmids. This approach provided insights into the role of microbial pathways in MASLD progression and highlighted key genetic signatures in both healthy and MASLD patients.

## 2. Results

### Agathobacter rectalis is depleted in MASLD

Certain bacterial clades have been positively correlated with MASLD progression, including *Enterobacteriaceae*^15,17^ such as *Escherichia / Shigella spp*.^17,18^, *Bacteroidaceae* like *Bacteroides spp.*^18,19^, *Veillonellaceae*^15^ or *Streptococcus spp.*^20^. Conversely, other clades have been reported to exhibit an inverse correlation with MASLD progression, including *Ruminococcaceae*^15^ such as *Faecalibacterium prausnitzii*^18,21^, *Lachnospiraceae* like *Eubacterium rectale*^18,21^ or *Dorea longicatena*^21^, *Clostridiaceae* like *Clostridium spp.*^17^, and even *Pseudomonas spp.*^20^. However, these studies often yield conflicting results. For example, a reported a positive correlation between *Ruminoccocus spp.* abundance and MASLD progression^19^ contradicts one of the main findings in two posterior studies^18,21^. Similarly, *Prevotella spp.* was found to decrease with MASLD development^22^ and fibrosis stage^19^, yet another study reported an increase in this genus in MASLD patients^20^. There are additional examples of discordant taxonomic signatures in MASLD, such as *Blautia spp.* or *Roseburia spp.*^23^.

To investigate whether any bacterial genera is systematically altered in MASLD, we profiled the taxonomic composition of the GM communities across all samples from the three cohorts at both genus and species level using MetaPhlAn^24^, as detailed in Materials and Methods. This analysis identified 28 bacterial genera shared across all three cohorts (Fig. 1A), which accounted for 59%, 31.5% and 49% of the GM fraction in each cohort, respectively (Fig. 1B). Thus, nearly half of the bacterial genera identified in at least one cohort were shared across all three (28 out of 63), representing between one-third and two-thirds of the total GM abundance.

**Figure 1.**
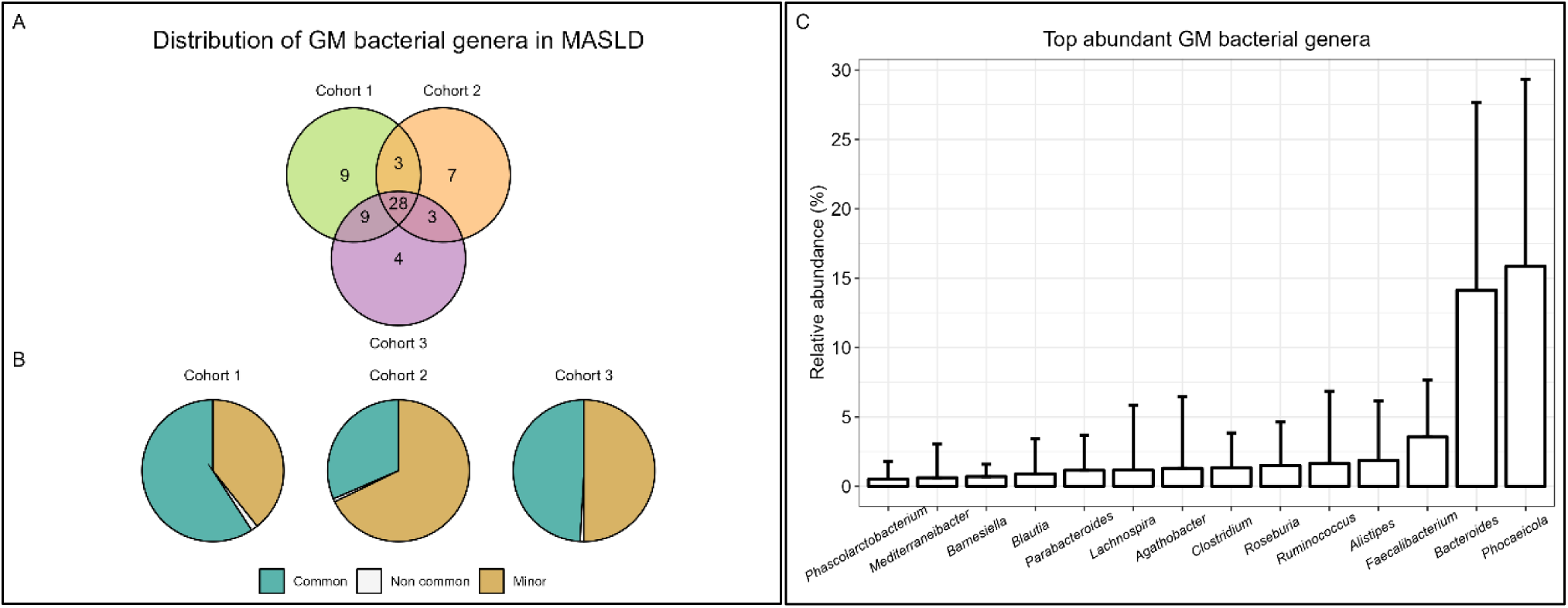
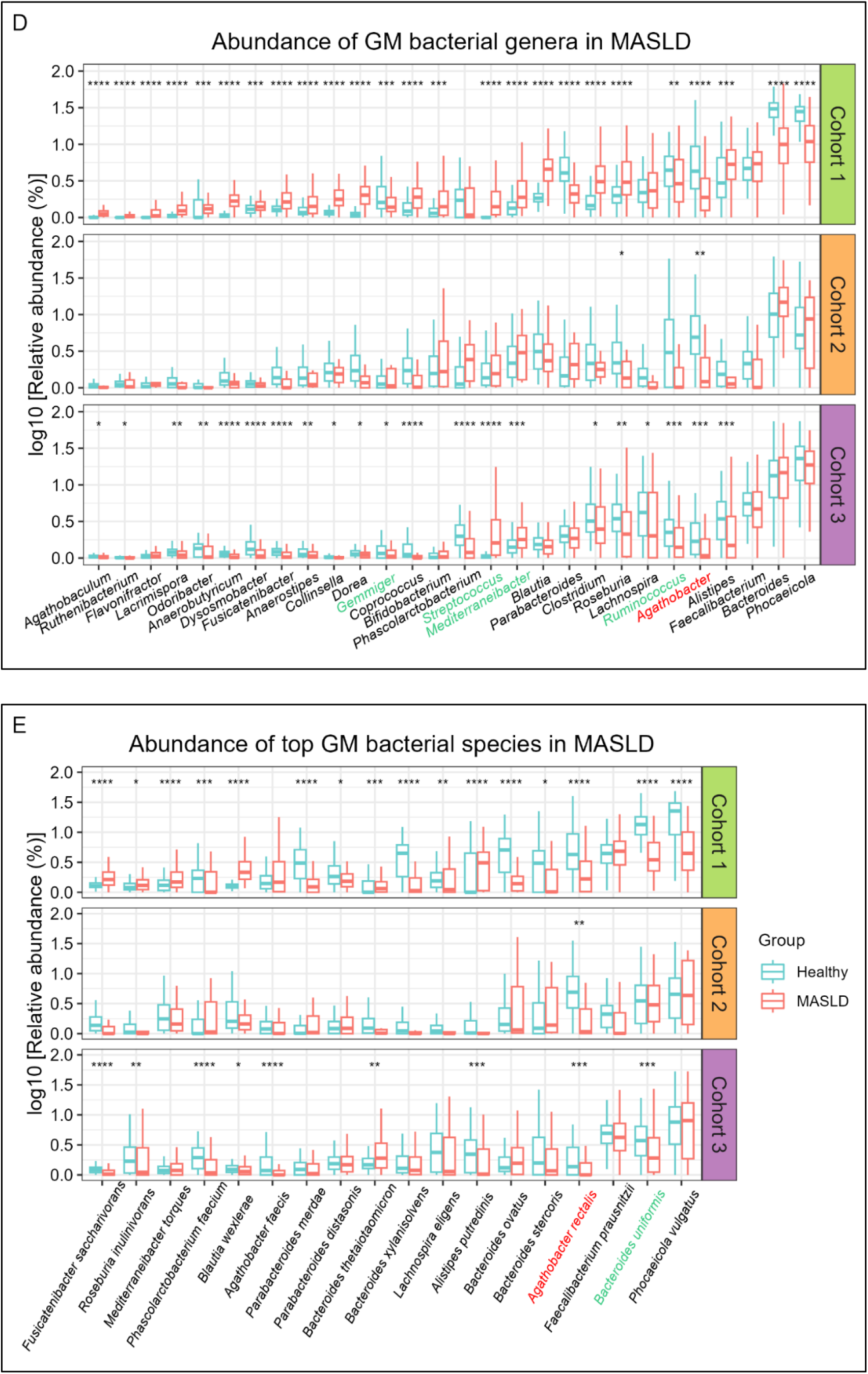
Taxonomic profiling of GM in MASLD. **(A)** Number of bacterial genera common to the three cohorts. **(B)** Average relative abundance of bacterial genera predicted by MetaPhlAn across the three study cohorts. The blue section of the pie charts reflects the average relative abundance of bacterial genera common to the three cohorts (n=28), whereas the grey and green sections reflect, respectively, the abundances of non-common and minor genera (i.e., with an average relative abundance of 0 in at least one of the cohorts). **(C)** Top abundant GM genera (average absolute abundance >0.5%) **(D-E)** Relative abundance of bacterial genera **(D)** and species **(E)** in the metagenomes of the fecal samples, shown on a logarithmic scale. Blue and red boxplots correspond to healthy and MASLD groups, respectively. Each facet represents one cohort. Differences in abundance were evaluated using pairwise Mann-Whitney tests with Benjamini-Hochberg adjustment for multiple testing. Horizontal bars represent median values, while boxes and whiskers represent the interquartile range and the full data spread (minimum to maximum), respectively. Clades depicted in green across the x-axis have a consistent differential abundance pattern between groups over the three cohorts. Only *Agathobacter rectalis*, depicted in red, has a consistent and statistically significant decrease in the MASLD group across cohorts. (*) p<0.05, (**) p<0.01, (***) p<0.001, (****) p<0.0001. Non-starred genera present non-significant differences in abundance between groups.

Among these shared genera, *Gemmiger*, *Streptococcus*, *Mediterraneibacter*, *Ruminococcus* and *Agathobacter* exhibited a consistent pattern of differential abundance between groups across all cohorts, reaching statistical significance in at least two (Fig. 1D). This exclusion criterion was applied to enhance the reliability of the results, ensuring that differentially abundant genera were not solely driven by heterogeneous demographic variables (e.g. geographical location, age, sex, or ethnicity) or environmental factors (e.g. diet, physical activity, comorbidities, or drug exposure), rather than being directly related to the MASLD phenotype^25,26^. However, this threshold was not applied to Cohort 2 because all patients in this group had MASLD, and applying it could have been overly restrictive, masking potential taxonomic differences associated with the disease. Of these five genera, only *Agathobacter spp.* showed statistically significant differences across the three cohorts (Fig. 1D). Note that *Agathobacter*, along with *Ruminococcocus* and *Mediterraneibacter*, belongs to the fraction of the highest abundant GM bacterial genera (Fig. 1C), highlighting their potential contribution to the phenotype in specific patient cohorts.

Analogously, the most abundant bacterial species within the GM were analyzed for differences in abundances between groups across the three cohorts (Fig. 1E). Only *Agathobacter rectalis* and *Bacteroides uniformis*, two of the most abundant GM bacterial species (Supplementary Fig. 1), exhibited a consistent pattern of differential abundance across all cohorts, with statistical significance in at least two. However, only *A. rectalis* showed a significant difference between groups in all cohorts, identifying it as a robust taxonomic marker depleted in MASLD.

### Butyrate-producing genes are depleted in MASLD

The identification of *A. rectalis* as a consistent taxonomic signature across multiple MASLD cohorts underscores its potential protective role in MASLD development. However, taxonomy alone may not fully capture the functional contributions of the GM. Certain microbial metabolites regulate GLA integrity and have been suggested to influence MASLD progression^27–29^. These compounds derive from dietary carbohydrates and proteins that are not fully digested in the upper gastrointestinal tract and are instead catabolized by the GM through fermentative pathways that are predominant in the anaerobic environment of the colon^30,31^. Once absorbed into the bloodstream, they are transported to the liver via the portal vein, where they can exert both beneficial and detrimental effects^32,33^.

The most well-characterized of these metabolites are short chain fatty acids, produced by the colonic fermentation of complex carbohydrates present in dietary fiber that escape digestion in the small intestine. These metabolic pathways have been relatively well understood for nearly thirty years^34^ and their impact on GM homeostasis has been reviewed in several articles^35–37^. For this reason, we interrogated the metagenomic fecal samples from the patient cohorts for metabolic signatures that could explain the MASLD phenotype. First, we focused on the genes encoding enzymes involved in butyrate formation as a proxy for the bacteria that carry them, independently of their taxa. To validate our hypothesis, we analyzed the genic abundance of the final enzymes involved in butyrate production. We selected butyrate as a target because it is crucial for maintaining GLA homeostasis by supporting the colonic epithelial barrier^38,39^. It also promotes lipid oxidation^40,41^, induces hepatic gluconeogenesis^42^, and prevents insulin resistance^43^, all of which are key factors in MASLD. In mice, both butyrate-producing probiotics^44^ and sodium butyrate administration^45–47^ have been shown to prevent steatosis and slow MASLD progression. Additionally, fecal microbiota transplantation (FMT) from healthy to MASH mice increased cecal butyrate levels and reduced steatosis^48^.

Bacterial butyrate can be formed through two major terminal reactions (Fig. 2A). The first involves a dephosphorylation of butyryl-P that yields one ATP through an energetically favorable reaction that is catalyzed by a butyrate kinase encoded by *buk.* In the precursor reaction, butyryl-P is formed through a phosphorylation of butyryl-CoA catalyzed by a phosphate butyryl transferase encoded by *ptb*. Butyrate also accumulates after the transfer of a CoA cofactor from butyryl-CoA to acetate through butyryl-CoA:acetate CoA transferase, encoded by *but*. This CoA-transferase pathway also conserves the energy of the CoA bond in the newly formed CoA-moiety of the co-substrate. Butyryl-CoA is produced from crotonyl-CoA through two different enzymes: butyryl-CoA trans-2-oxidorreductase, encoded by *fabV*, and butyryl-CoA dehydrogenase, encoded by *bcd.* This pathway is further reviewed in the human GM elsewhere^49^.

**Figure 2.**
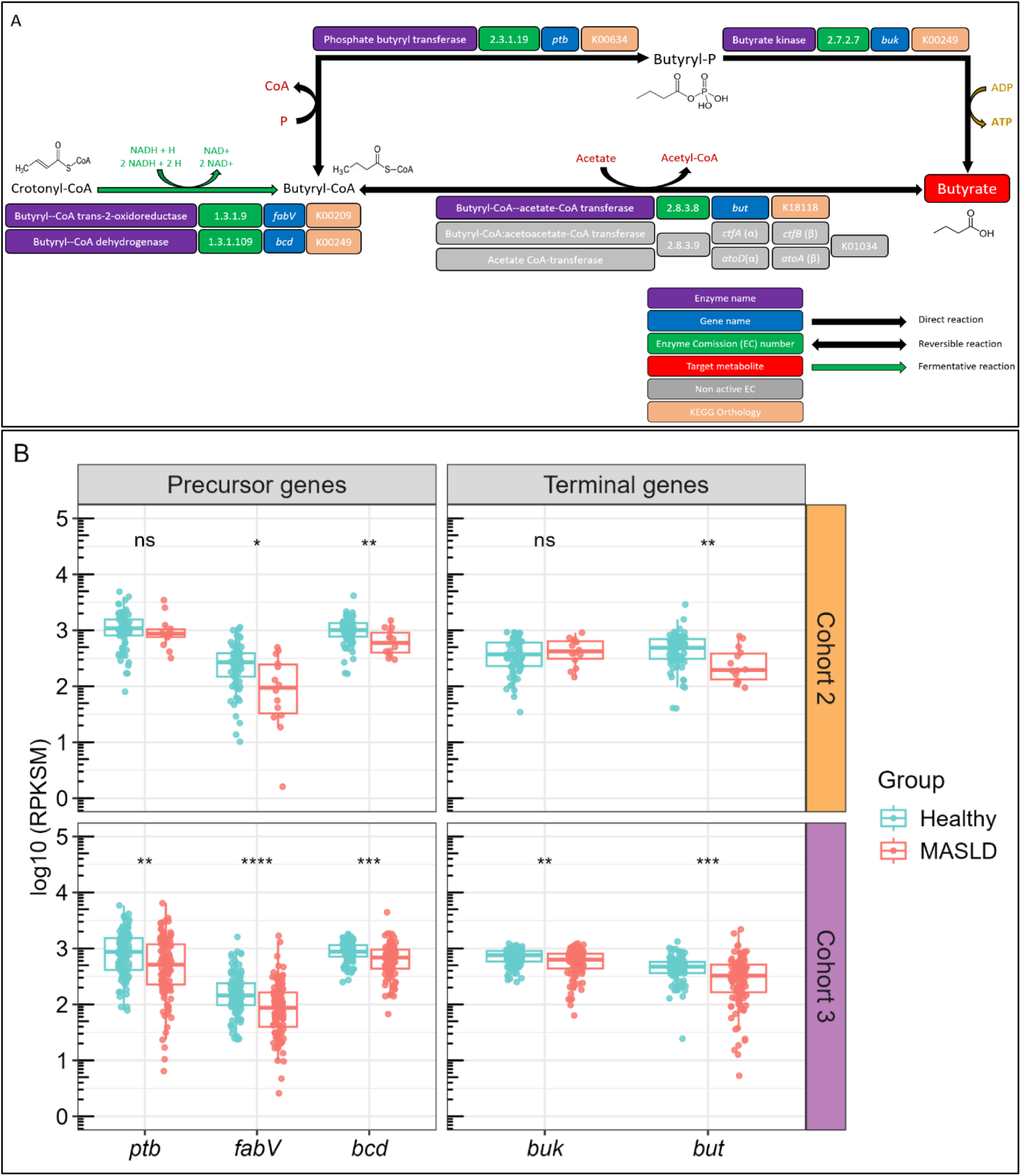
Abundance of butyrate-producing genes in patient cohorts. **(A)** Characterization of the terminal reactions involved in the formation of bacterial butyrate from crotonyl-CoA. Enzymes, EC numbers, coding genes and KOs, as well as reaction types, are indicated for each metabolic step according to the legend (lower part). **(B)** Abundance of butyrate-producing genes in Cohorts 2 and 3, shown on a logarithmic scale. Boxplots represent the gene abundance from the metagenomic measures of the fecal samples, and differences in abundances were evaluated using pairwise Mann-Whitney tests with Benjamini-Hochberg adjustment for multiple testing. Upper and lower facets represent abundances in Cohorts 2 and 3, respectively. Left and right facets represent abundances of precursor and terminal coding genes, respectively, as described in (A). RPKSM: reads per kilobase per genic size per million reads. (*) p<0.05, (**) p<0.01, (***) p<0.001, (****) p<0.0001, ns: non-significant.

We quantified the abundance of these genes in the metagenomic fecal samples of both groups across the three MASLD cohorts, as described in Materials and Methods. Genes involved in the production of butyrate and its precursors, butyryl-P and butyryl-CoA, were significantly depleted in the GM of patients with cirrhosis MASLD compared to the healthy group, and their abundance also decreased significantly with MASLD progression (Fig. 2B). The precursor gene *fabV* was nearly an order of magnitude less abundant than *ptb* and *bcd* in both cohorts, underscoring the greater prevalence of *bcd* over *fabV* in the CoA-transferase route. Additionally, *ptb* did not exhibit significant differences in Cohort 2. Gene *but* was significantly depleted in the control group in both cohorts, whereas *buk* did not show differences in Cohort 2 either. Notably, the depletion of butyrate-related genes was not observed in MASLD patients from Cohort 1.

### SCA-producing genes are increased in MASLD

Short-chain alcohols (SCAs) are a metabolic family also suggested to regulate GLA integrity, with several studies linking GM endogenous alcohol production to MASLD. One of the first reports, published over a decade ago, described elevated peripheral blood ethanol and increased fecal *Escherichia spp.* 16S rRNA levels in patients with MASH^50^. Similar findings were reported in a pediatric cohort, where children with MASLD exhibited higher levels of fecal ethanol, *Prevotella spp.*, and Gammaproteobacteria 16S rRNA^51^. A more recent study showed that postprandial peripheral blood ethanol and fecal *Lactobacillaceae* 16S rRNA levels increased in MASH patients after selective inhibition of host ADH. This effect was reversed with broad-spectrum antibiotic treatment^52^.

In 2019, Yuan and colleagues identified a high-alcohol-producing strain of *Klebsiella pneumoniae* capable of causing MASLD through ethanol production^53^. Specifically, they found that FMT from a MASH patient containing this strain induced MASLD in healthy mice, whereas removing the strain before FMT prevented disease development. Furthermore, ethanol produced by this *K. pneumoniae* strain was shown to disrupt lipid homeostasis in hepatic cells and to cause mitochondrial dysfunction, oxidative stress, and the accumulation of reactive oxygen species^54^. These factor are critical to MASH development through their role in lipid peroxidation and inflammation^55^. Additionally, a recent computational study identified a positive correlation between the abundance of two fermentative pathways in certain families from class Clostridia and fatty-liver disease in patients. These pathways include the production of ethanol from pyruvate and the heterolactic fermentative pathway^56^.

With this ample evidence, we hypothesized that genes encoding enzymes involved in the formation of endogenous SCAs may represent another metabolic mechanism contributing to MASLD pathology. We focused on ethanol and propanol, which are produced by the reduction of their respective aldehydes through ethanol and propanol dehydrogenases. These enzymes help to reduce the levels of toxic aldehydes in the GM^57,58^ and replenish NAD^+^ levels to maintain fermentative growth. Bacterial ethanol is formed from acetaldehyde through a fermentative reaction that can be catalyzed by 1) an NADH-dependent, bifunctional acetaldehyde-ethanol dehydrogenase encoded by *adhE*, 2) an NADPH-dependent alcohol dehydrogenase encoded by *adhB*^59,60^ and 3) a NADPH-dependent aldehyde reductase encoded by two different genes: *ahr* and *yahK*, with the corresponding liberation of phosphorylated reducing power in form of NADP^+^ ^61,62^. The precursor acetaldehyde can be formed from 1) ethanolamine degradation, through a reaction catalyzed by ethanolamine ammonia-lyase which is encoded by *eutB* and *eutC* genes and liberates ammonia as a by-product^63^, 2) acetyl-CoA, through a fermentative reaction catalyzed by bifunctional AdhE and by a CoA-acetylating acetaldehyde dehydrogenase encoded by *mhpF*^64,65^ and 3) acetate, through a dehydration catalyzed by aldehyde dehydrogenase B, encoded by *aldB*^66^ (Fig. 3A, top). Bacterial 1-propanol is formed from propionaldehyde through a fermentative reaction catalyzed by a propanol dehydrogenase encoded by two different genes: *adhP* (a propanol-preferring alcohol dehydrogenase) and *pduQ.* Propionaldehyde can be generated through two distinct reactions: 1) a CoA transference from propionyl-CoA catalyzed by a propionaldehyde dehydrogenase encoded by *pduP*; and 2) a dehydration of 1,2 propanediol catalyzed by a propanediol dehydratase composed by three subunits encoded by *pduC*, *pduD* and *pduE.* These genes are co-located within the same operon and encode the corresponding enzymes responsible for the metabolism of 1,2-propanediol within the propanediol-utilizing (Pdu) microcompartment, a specialized bacterial structure found in certain enteric species^67,68^ (Fig. 3A, bottom).

**Figure 3.**
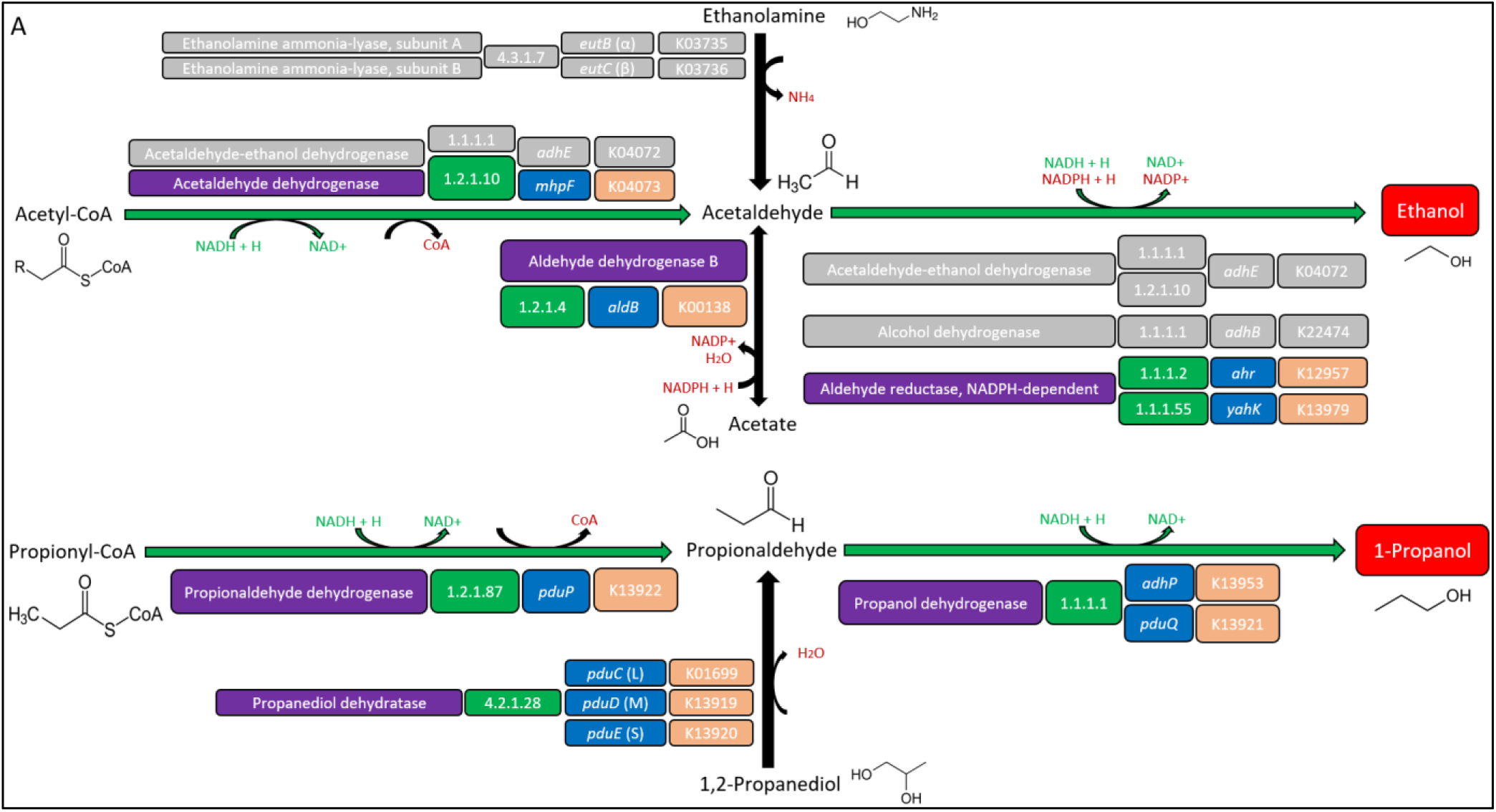
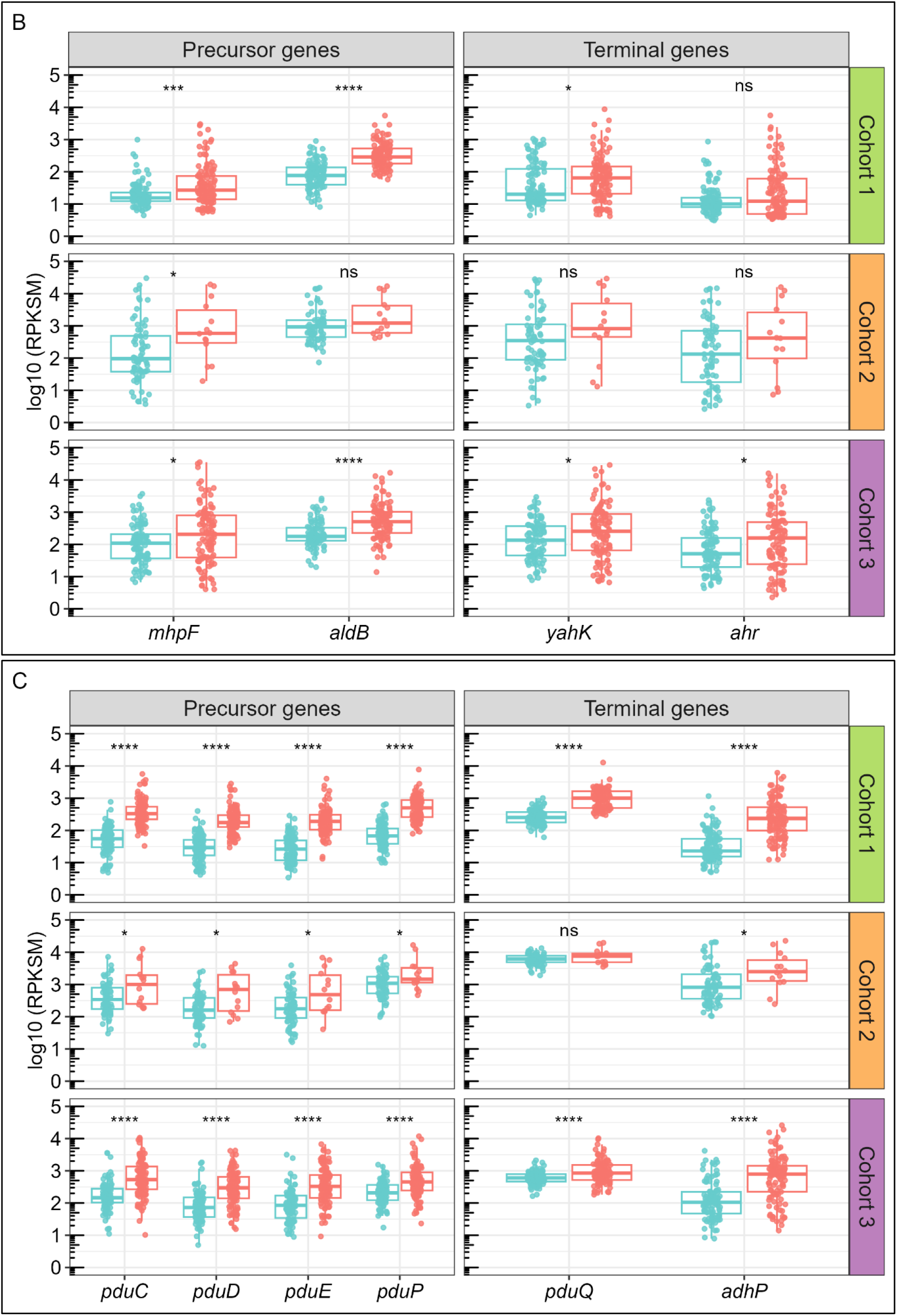
Abundance of SCA-producing genes in patient cohorts. **(A)** Characterization of the terminal reactions involved in the formation of bacterial ethanol (top) and propanol (bottom). Enzymes, EC numbers, coding genes and KOs, as well as reaction types, are indicated for each metabolic step according to the legend in Fig. 2A. Non-candidate genes are colored in grey. **(B)** Abundance of *mhpF, aldB, yahK* and *ahr* genes involved in the last two reactions from pyruvate to ethanol in the three cohorts. **(C)** Abundance of *pduC-E, pduP, pduQ* and *adhP* genes, involved in the last two reactions from 1,2-propanediol and propionyl-CoA to 1-propanol in the three cohorts. Boxplot layout, distribution in facets and statistical tests were performed as in Fig. 2C.

We quantified the abundance of genes involved in bacterial SCA formation in the metagenomic samples of the three cohorts as described above. The genes *mhpF* and *aldB*, involved in ethanol production, and *yahK* and *ahr*, involved in acetaldehyde production, were significantly enriched in the GM of MASLD patients from Cohort 1 and Cohort 3 compared to healthy groups. Their abundance also increased with MASLD progression (Fig. 3B). Notably, *mhpF* and *ahr* were depleted compared to *aldB* and *yahK* respectively in all three cohorts. All genes involved in the formation of 1-propanol and propionaldehyde were also significantly enriched in MASLD patients by more than half order of magnitude, increasing with MASLD progression (Fig. 3C, Supplementary Fig. 4A). These findings reveal a systematic increase of SCA-producing genes in MASLD.

### Genes involved in methane production are decreased in MASLD, whereas *tor* operons driving TMA accumulation are elevated

Trimethylamine N-oxide (TMAO) is a metabolite derived from the oxidation of trimethylamine (TMA), which is produced by the microbial degradation of dietary compounds such as choline and L-carnitine. Circulating TMAO levels have also been linked to MASLD severity^69^ and all-cause mortality in MASLD patients^70^. Higher plasmatic TMAO levels also increase cardiovascular disease risk-a condition associated with MASLD-in both mice^71^ and clinical patients^72^. Therefore, we investigated whether bacterial TMAO could represent another metabolic mechanism contributing to MASLD.

Methane production in the GM occurs through a two-step process: a simultaneous TMA demethylation and CoM methylation, followed by the reduction of methyl-CoM (Fig. 4A). TMA originates from choline degradation via the *cut* operon^73^ and is oxidized in the liver to TMAO^74^ or reduced back to TMA under anaerobic conditions via the *torCAD* and *torZY* operons. Specifically, *torA* and *torZ* encode two TMAO reductases, whose molecular mechanisms are reviewed elsewhere^75^.

**Figure 4.**
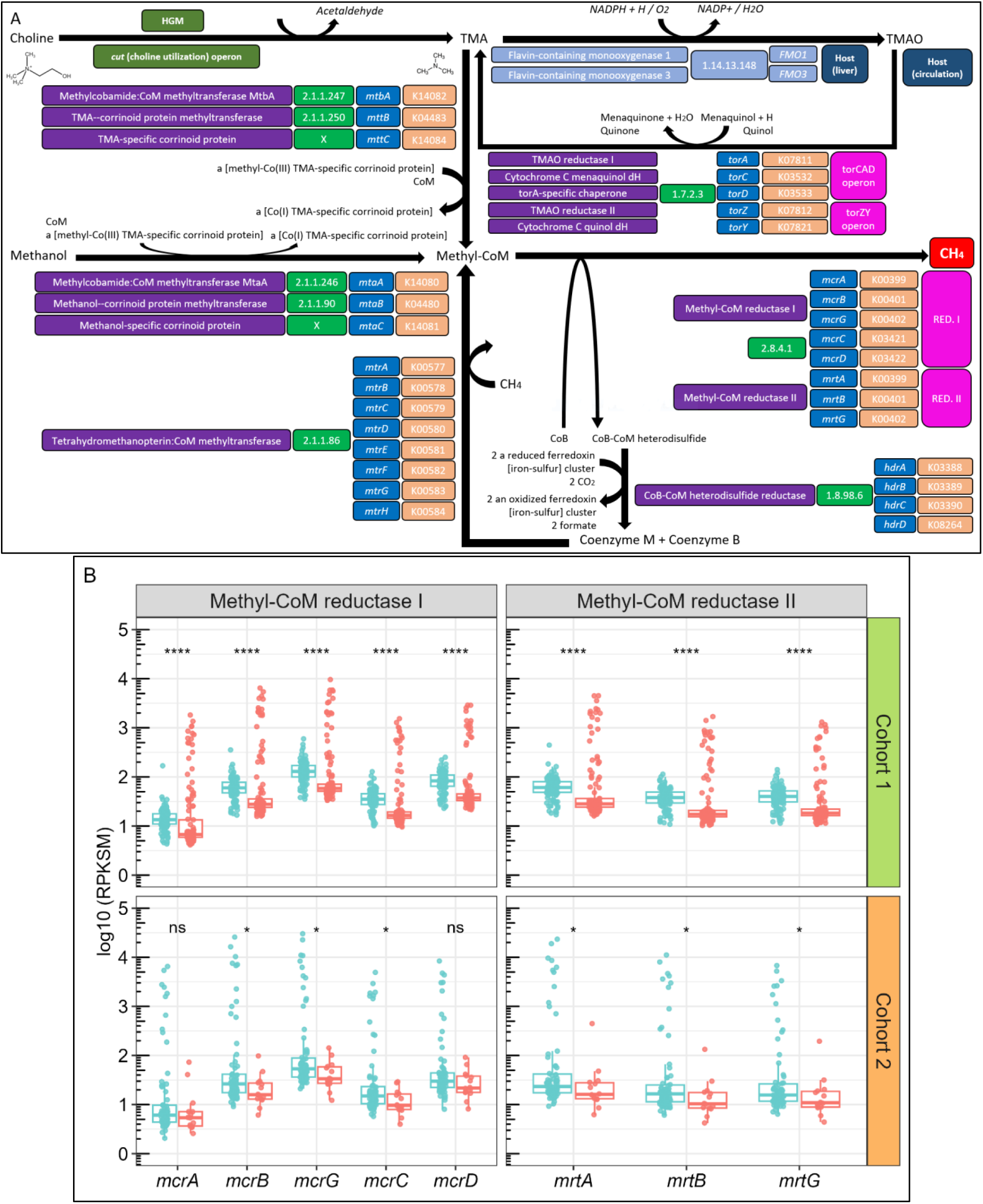
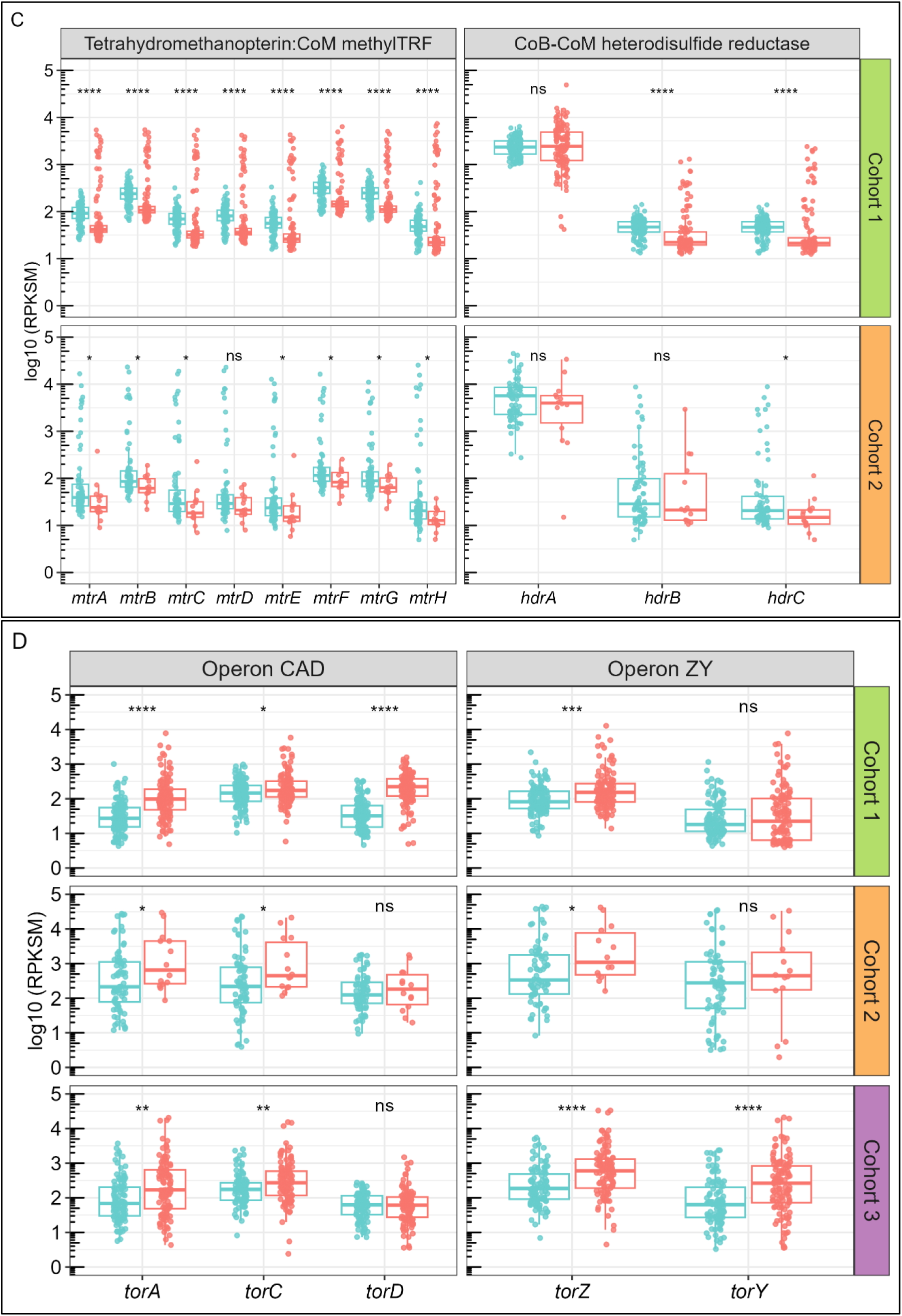
Abundance of methane and TMA-producing genes in patient cohorts. **(A)** Characterization of the terminal reactions involved in the formation of microbial methane. Enzymes, EC numbers, coding genes and KOs, as well as reaction types, are indicated for each metabolic step according to the legend annexed to Fig. 2A. **(B)** Abundance of genes encoding methyl-coM reductase I (*mcr*) and methyl-CoM reductase II (*mrt*). **(C)** Abundance of genes encoding CoB-CoM heterodisulfide reductase (*hdr*) and tetrahydromethanopterin S-methyltransferase (*mtr*). **(D)** Abundance of *tor* operons. Facets represent genic operons. Boxplot organization and statistical tests were conducted as in Fig. 2C.

CoM methylation occurs through two well-characterized enzymatic systems: a TMA-specific (composed of *mttC*, *mtbA* and *mttB*) and a methanol-specific pathway (composed of *mtaC, mtaA* and *mtaB*)^76^. The final reduction of methyl-CoM to methane is catalyzed by methyl-CoM reductase (MCR) I, which is encoded by the *mcrBDCGA* operon^77^. MCR I has an alternative isozyme termed MCR II, encoded by *mrtBGA* operon^78^. The resulting heterodisulfide (CoB-S-S-CoM) is then reduced to CoM and CoB by a heterodisulfide reductase encoded by operon *hdrABC*, thereby regenerating both cofactors^79^. Additionally, methyl-CoM is replenished via tetrahydromethanopterin CoM-methyltransferase (encoded by operon *mtrABCDEFGH*), which methylates CoM^80^.

We measured the abundance of these genes in the samples of the three MASLD cohorts as described above. Genes encoding MCR I and II, which catalyze the reduction of methyl-CoM to methane, were significantly decreased in the GM of MASLD patients. Their abundance also declined with MASLD progression (Fig. 4B). Concordantly, *hdr* and *mtr* genes, responsible for replenishing CoM and transferring it to a methyl group to form the methyl-CoM precursor, were significantly decreased in MASLD and showed a progressive depletion with MASLD advancement (Fig. 4C). Interestingly, *hdrA* was two orders of magnitude more abundant than the rest of the genes from the operon. Genes *mtbA, mttB, mtaA* and *mtaB* were significantly decreased in MASLD. However, they did not exhibit differences between the initial and advanced stages of MASLD (Supplementary Fig. 2). Notably, the higher abundance of methane-producing genes observed in the healthy group was not replicated in Cohort 3. In contrast, bacterial genes involved in TMA regeneration showed the opposite trend. The *torCAD* and *torZY* operons were significantly increased in the GM of MASLD patients, and their abundance increased with MASLD progression (Fig. 4D, Supplementary Fig. 4B). These findings suggest that methane production from TMA and methanol, which typically occurs in a healthy state, is disrupted in MASLD, revealing another potential microbial pathway involved in MASLD pathogenesis.

Finally, we quantified the abundance of five universal single-copy genes (USCGs), as described in Material and Methods, to confirm that observed differences in genic abundance between sample groups were truly associated with the MASLD phenotype and not confounded by biological variability or spurious technical artifacts related to the quantification method. No significant differences were found in gene abundance between groups for any USCG in any of the three cohorts (Supplementary Fig. 1).

### Candidate metabolic genes are accessory in the GM

Although this analysis identifies *A. rectalis* and multiple metabolic genes involved in the formation of butyrate, SCAs, TMAO and methane as consistent taxonomic and functional signatures associated with MASLD, these markers alone are not sufficiently robust to establish a causal link between GM composition and MASLD phenotype. GM bacterial species have pleomorphic genomes, consisting of up to thousands of genomically distinct strains that share a conserved genetic core but possess an accessory genome that is highly variable among them^81,82^. This accessory genome is often encoded in plasmids and other MGEs which are subject to frequent changes and can readily transfer between different bacterial strains^16,83^.

To test this hypothesis, we assessed whether candidate metabolic genes involved in MASLD are core or accessory across some of the most abundant human GM genomes. Genome selection and genic presence were determined as described in Materials and Methods. Briefly, the presence of a target gene in a clade was defined as the percentage of strains encoding it, classifying it as core if present in >80% of congeneric or conspecific genomes, as accessory if found in 20-80%, and as highly accessory if present in <20% genomes.

Most of the selected GM genera (n=24) belong to class Clostridia, specifically to orders Lachnospirales and Oscillospirales (Fig. 5A). Butyrate-producing genes from the butyryl-CoA pathway (*bcd* and *but*) were core in nearly half of these genera, while those from the butyryl-P pathway (*ptb* and *buk*) were core only in *Flavonifractor, Clostridium* and *Coprococcus.* Notably, butyrate-producing genes were highly accessory across several genera, including *Lachnospira, Ruminococcus_B* and *E, Ruthenibacterium, Blautia*, *Fusicatenibacter*, *Dorea,* and *Mediterraneibacter*, and were completely absent in *Ruminococcus_C* and *D*. In contrast, acetaldehyde and ethanol-producing genes were highly accessory across all genera except *Streptococcus*, an heterolactic genus that constitutively encoded *aldhB* in all its genomes (specifically in *S. salivarius* and *S. thermophilus,* Fig. 5B) and exhibited moderated presence (20-80%) of *yahK*/*ahr*. The acetaldehyde-producing gene *mhpF* was nearly absent in the GM, except for a few genomes of *Flavonifractor* and *Agathobacter*. Propionaldehyde and propanol-producing genes from the *pdu* operon were core in three genera: *Ruminococcus_B* (*R_B. gnavus*), *Anaerobutyricum* (*A. hallii*) and *Flavonifractor* (*F. plautii*), though *pduE* was unexpectedly absent in *Ruminococcus_B*. These genes were highly accessory in *Blautia* (*B. wexlerae*), *Streptococcus*, *Medierraneibacter* or *Agathobacter* (*A. rectalis*). Notably, *A. hallii* and *F. plautii* were the only species encoding both core butyrate and propanol-producing genes.

**Figure 5.**
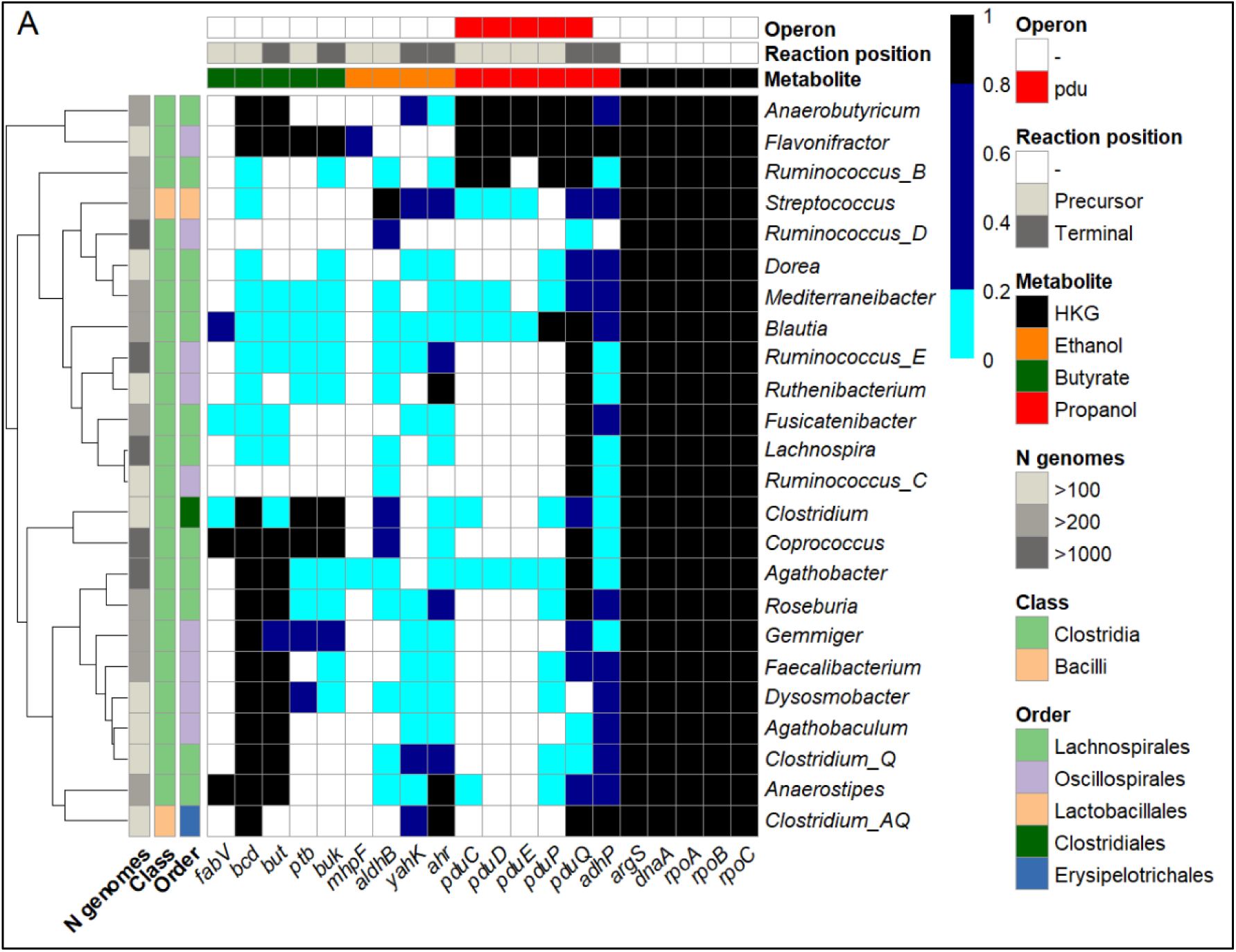
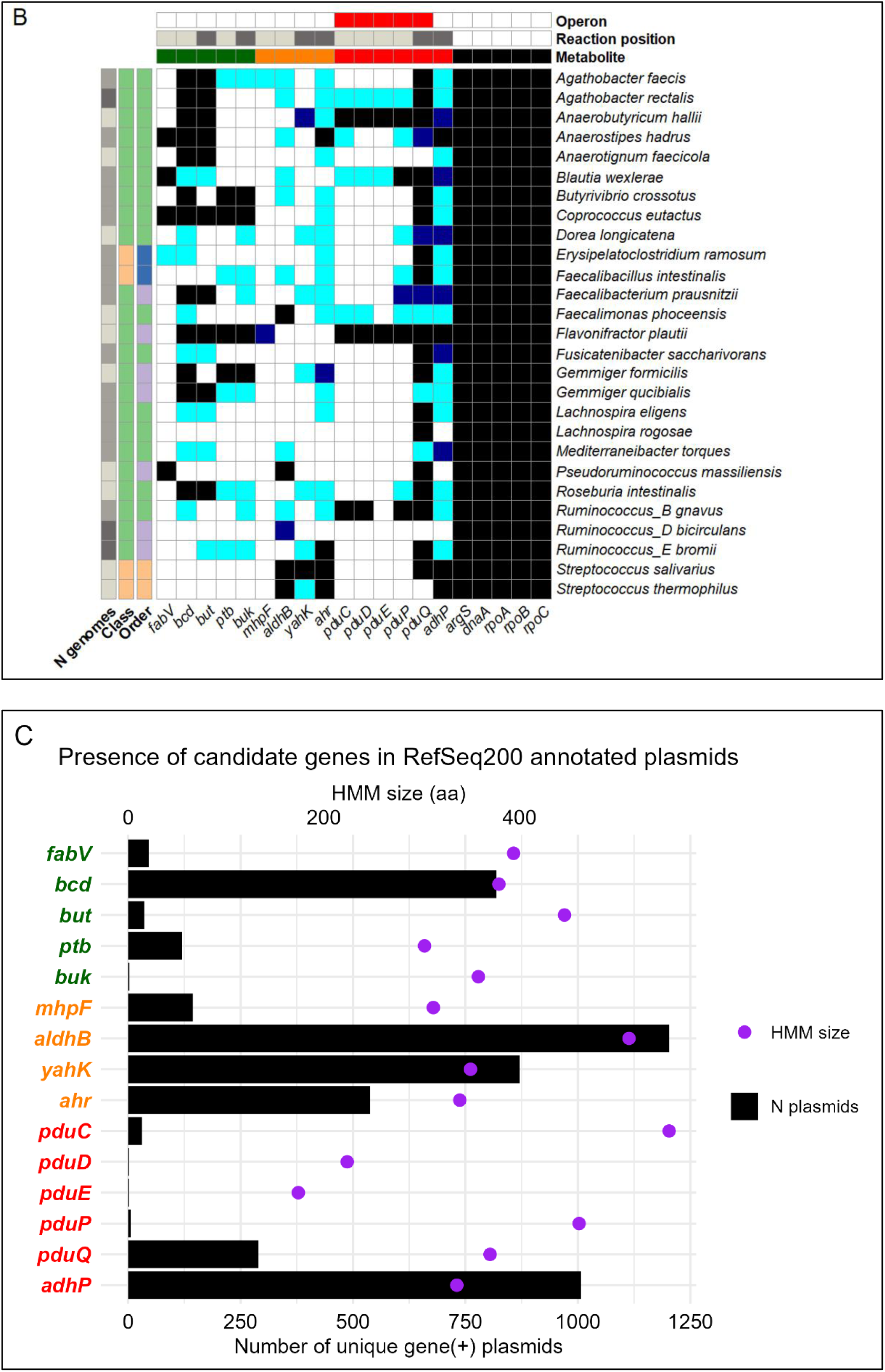
Presence profile of candidate metabolic genes in the human GM. **(A-B)** Heatmap showing the presence of candidate genes involved in MASLD across some of the most abundant clades in the human GM. The x-axis represents genes associated with butyrate and SCAs production, as well as USCGs. The y-axis lists the bacterial clades inspected. Gene presence is expressed as the percentage of **(A)** congeneric and **(B)** conspecific genomes encoding each target gene, with colors indicating classification: black for core genes (>80% of genomes), dark blue for accessory genes (20-80% genomes), light blue for highly accessory genes (<20% genomes) and white for genes completely absent in the clade. Hierarchical clustering was applied to the y-axis to group genera (top) based on gene-presence similarity. **(C)** Presence of candidate genes in RefSeq200 annotated plasmids. Black bars represent the number of different plasmids that encode at least one copy of each gene.

Note that the nomenclature used in Fig. 5 follows the Genome Taxonomy Database standards, which redefine taxonomic ranks based on genome-wide phylogenetic analyses^84^. As a result, traditional genera such as *Ruminococcus* and *Clostridium*, which were found to be polyphyletic (i.e., composed of species that do not share a single common ancestor) are split into multiple monophyletic clades (e.g., *Ruminococcus_A, Ruminococcus_B* or *ClostridiumQ*).

Beyond their accessory nature within GM pangenomes, candidate metabolic genes were also encoded on plasmids (Fig. 5C). Genes involved in ethanol formation such as *aldhB, yahK* and *ahr* were among the most widely distributed, being present in hundreds of different plasmids. In contrast, butyrate-producing genes (*fabV, but, ptb,* and *buk*) and propanol-related genes (*pduC, pduD, pduE, pduP,* and *pduQ*) were detected in fewer plasmids, further supporting their variable presence across GM genomes. A notable exception was *bcd*, which exhibited a broader presence across both chromosomal and plasmid-encoded sequences. These findings reinforce the idea that non-essential metabolic functions, especially ethanol production, are encoded in MGEs, contributing to their accessory nature within the GM.

## 3. Discussion

MASLD is a liver pathology influenced by multiple contributing factors, and the GM has been long suspected to play a role in its progression. Numerous studies have tried to ascribe a causality between GM composition and MASLD, but taxonomic signatures in key GM clades, such as *Blautia* and *Roseburia spp.*, have been inconsistently associated with MASLD progression^23^. These issues may be linked to the use of 16S rRNA-based profiling in metataxonomic studies, which frequently lacks resolution at species and strain level^85^.

However, the heterogeneity concomitant to bacterial genomes poses another possible cause for the lack of association between disease progression and the taxonomic distribution. Bacterial genomes are highly plastic, and strains from the same species often harbor different genes, many of which are involved in different metabolic pathways^81^. Fermentative pathways in lactobacilli, for example, are often strain-specific^86^, and the products of the enterobacterial mixed acid fermentation also exhibit wide variability between strains^87^. If MASLD progression is linked to the metabolic functions of the GM, then the association between the disease and the taxonomic profile may be obscured by the inherent metabolic diversity of bacterial species. To study these potential alternatives, we followed two distinct strategies.

To identify species associated with MASLD and overcome the limitations of 16S rRNA analysis, we utilized multiple marker genes for taxonomic classification, in an attempt to provide a more comprehensive view of microbial abundances in health and disease states. Using MetaPhlan, we profiled the taxonomic composition of the GM in metagenomic fecal samples of healthy and MASLD donors across three independent cohorts. We identified 28 bacterial genera shared across all cohorts (Fig. 1A), collectively accounting for 30-60% of the total GM abundance (Fig. 1B). Among these, five candidate genera-*Gemmiger*, *Streptococcus*, *Mediterraneibacter*, *Ruminococcus* and *Agathobacter* (Fig. 1C)- and two species, *Agathobacter rectalis* and *Bacteroides uniformis* (Fig. 1E), exhibited consistent differences in their relative abundance between groups across all the cohorts. These taxa rank among the most abundant clades in the GM (Fig. 1C, Supplementary Fig. 1). However, only *Agathobacter rectalis* showed statistically significant differences. Given its high abundance in the GM, its consistent abundance trend at both genus and species level, and its significant depletion in MASLD, *A. rectalis* emerges as a robust taxonomic marker for the disease.

Although this observation has been previously reported^18,21^, this result should be interpreted with caution due to ongoing debates in bacterial nomenclature, as *A. rectalis* is a newer designation that has replaced *Eubacterium rectale*^88^. Updates in taxonomic assignments pose challenges, as newer species or genera may not be consistently reflected across different databases or studies^84,89^. Moreover, taxonomic classifications themselves, whether based solely on 16S rRNA or on multiple markers, limit our ability to reliably infer GM changes. Bacterial genome plasticity, driven by MGEs, can blur the lines of taxonomy-phenotype associations, as they contribute to a vast strain-level diversity^81^ that remains undetected by taxonomic profiling methods. In fact, we have previously reported that geno-metabolic shifts in GM propionate production associated with inflammatory bowel disease do not align with species-level variations^90^.

GM associated metabolism contributes to MASLD by disrupting the integrity of the GLA. For this reason, we focused on identifying abundance differences in the genes responsible for producing metabolites potentially implicated in MASLD across several patient cohorts, i.e. butyrate, SCAs, TMA and methane. We isolated the GM gene families encoding the enzymes involved in the biosynthesis of these metabolites and aligned them against the metagenomic samples to quantify their abundance. We found that butyrate-producing genes are depleted in the GM of MASLD patients (Fig. 2B), particularly those involved in the crotonyl-butyryl CoA axis: *fabV* (butyryl-CoA oxidoreductase), *bcd* (butyryl-CoA dehydrogenase) and *but* (butyryl-CoA:acetate-CoA transferase) (Fig. 2A). While *ptb* and *buk* were also higher in both healthy subcohorts, their levels did not differ significantly between initial and advanced MASLD groups within Cohort 2. This suggests that CoA transfer may be the predominant pathway for butyrate production in the GM, as previously reported^91^. The subtler differences in *ptb* and *buk* in Cohort 2 may reflect a common MASLD phenotype across different fibrosis stages, potentially influenced by the smaller sample size relative to Cohort 3. Additionally, two other enzymes-structured as two-subunit systems-, can transfer CoA from different cofactors to butyrate to produce butyryl-CoA: butyryl-CoA:acetoacetate-CoA transferase (encoded by *atoD* and *atoA,* EC 2.8.3.9) and acetate CoA-transferase (encoded by *ctfA* and *ctfB,* EC 2.8.3.9) (Fig. 2A). These were not included in this study since they catalyze the reverse reaction, consuming butyrate to form butyryl-CoA^92^. Notably, the depletion of butyrate genes was not observed in MASLD patients from Cohort 1.

Contrarily, genes involved in the formation of ethanol and propanol through connected fermentative reactions are generally increased in MASLD (Fig. 3). Genes *mhpF* and *aldB*-encoding two aldehyde dehydrogenases that produce ethanol-, and *yahK* and *ahr*-encoding two NADPH-dependent aldehyde reductases that form acetaldehyde-are significantly increased in the GM of MASLD patients compared to the healthy groups (Fig. 3B). Genes involved in the production of 1-propanol structured in operon *pdu* are also increased in MASLD (Fig. 3C, Supplementary Fig. 4A). Specifically, *pduCDE*-encoding a propanediol dehydratase-, *pduP*-encoding a propionaldehyde dehydrogenase- and *pduQ* and *adhP*-encoding two distinct propanol dehydrogenases-. The pronounced differential abundance of these propanol-producing genes, organized in the *pdu* operon and *adhP*, which have been largely unexplored in the context of liver disease, suggests that 1-propanol produced in bacterial microcompartments may play a more significant role than ethanol in MASLD onset and progression.

The abundance of genes in the TMA-methane metabolic axis is also disrupted in MASLD. Genes involved in methane formation (Fig. 4A) are depleted in the GM of individuals with the pathology. This reduction includes both MCR I and II coding-genes (Fig. 4B), which reduce methyl-CoM to methane and *mtr*, *hdr* (Fig. 4C), and Mtb / Mta-coding genes (Supplementary Fig. 2), dedicated to regenerating methyl-CoM. In contrast, TMAO-reductive genes, organized in *torCAD* and *torZY* operons and responsible for regenerating TMA, are increased in MASLD (Fig. 3D, Supplementary Fig. 4B). These results suggest that methane production is impaired in MASLD. The opposing trend in the abundance of *tor* operons further indicates that this impairment stems from a metabolic bottleneck that leads to TMA accumulation instead of its conversion to methyl-CoM, effect that is driven by TMAO reductases I and II.

Taken together, these results reveal a geno-metabolic alteration associated with MASLD, characterized by an increase in genes involved in the production of SCAs and TMA, and a concurrent depletion of genes responsible for butyrate and methane production (Fig. 6). We confirmed that differences in gene abundance were driven by the MASLD phenotype by showing no differences in USCG abundance between groups (Supplementary Fig. 3). These findings support the hypothesis that the accumulation of endogenous alcohols may represent a key metabolic feature of MASLD, occurring alongside the displacement of butyrate-producing taxa, as previously proposed^33^. Moreover, the observed decrease in *A. rectalis* abundance partially explains the reduction in butyrate levels, as this organism is a known GM butyrate producer^88^.

**Figure 6.**
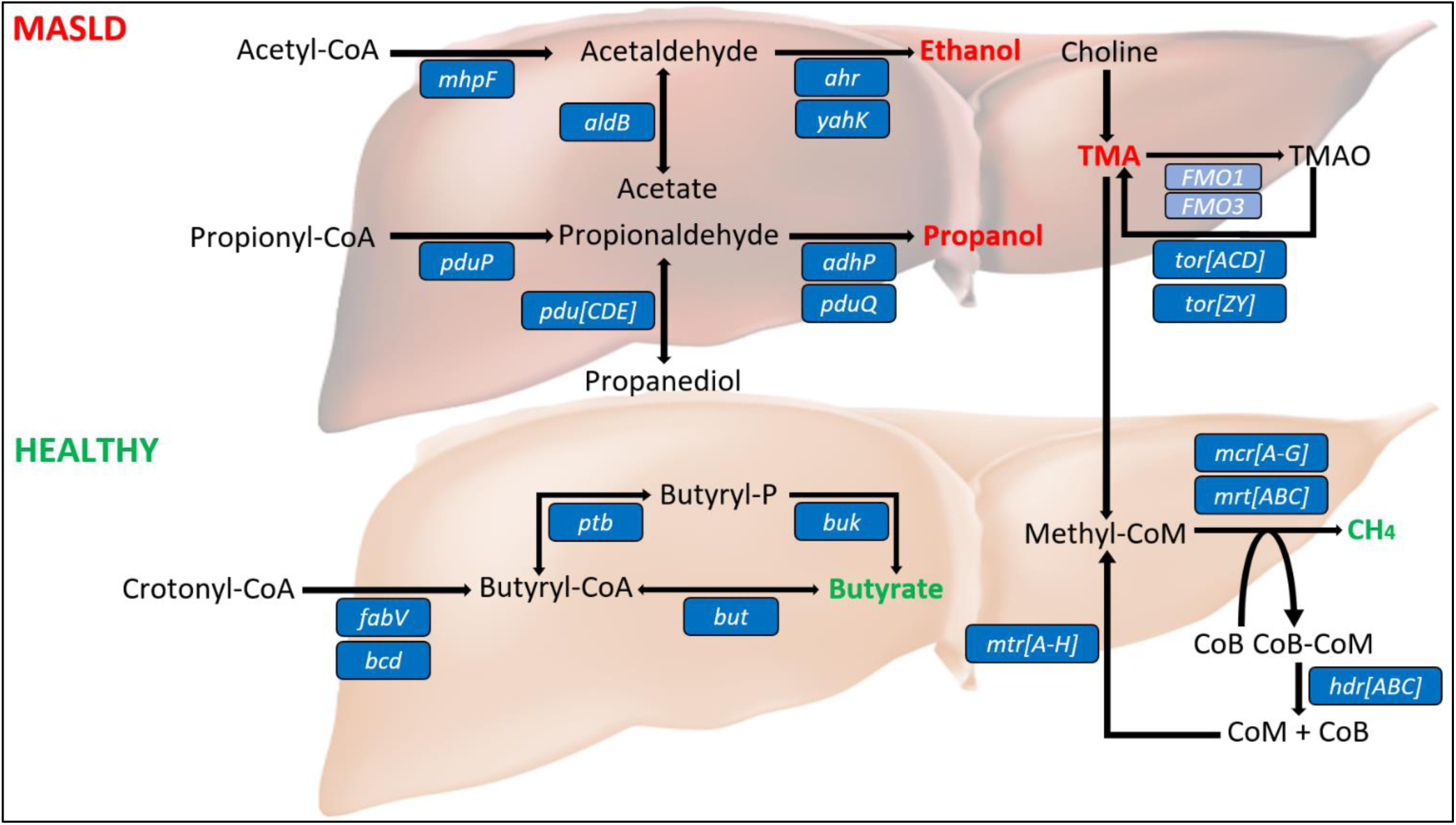
Geno-metabolic scheme involved in MASLD. Proposed scheme showing the GM genic alterations associated with butyrate, SCAs, TMA and methane production in MASLD. Genes involved in the production of ethanol, propanol and TMA are increased in the pathology (top), whereas those associated with butyrate and methane are decreased (bottom). Blue boxes indicate the catalyzing genes whose abundance is altered in each metabolic step.

**Figure 7.**
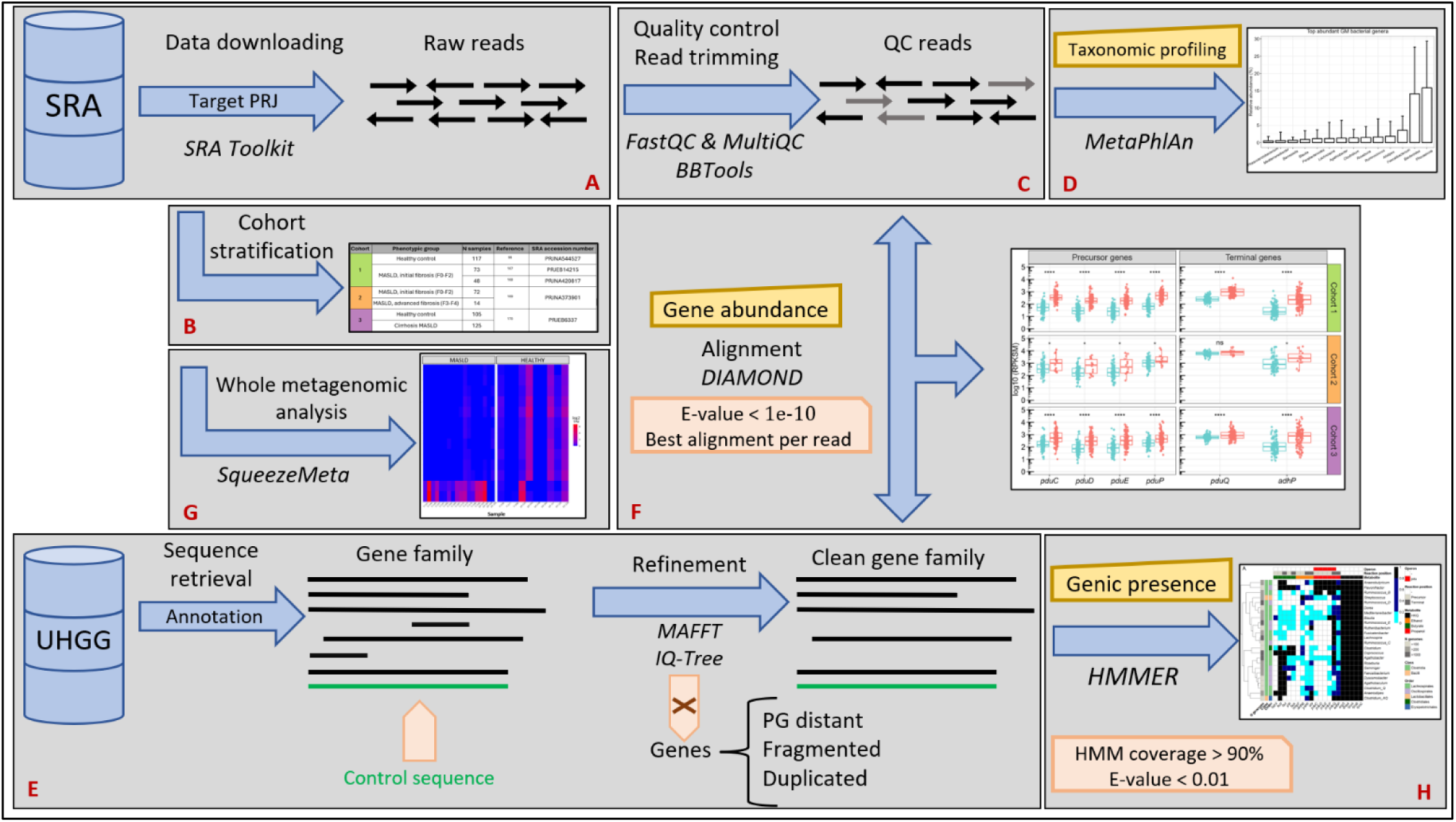
Overview of the methodology used in this study. This diagram summarizes the complete computational workflow for processing whole paired-end metagenomic samples, including data retrieval from the Short Read Archive (SRA), read quality control and taxonomic profiling. Target gene families were isolated and their abundance quantified by alignment against the sequencing libraries. Additionally, the presence of target genes was examined across GM genomes and annotated plasmids was inspected. Blue arrows indicate the progression between workflow steps, with the corresponding software shown in black beneath each arrow. Filtering steps are highlighted in pink boxes. Red-labeled references indicate the corresponding subsection in Materials and Methods (A-H), where each step is explained in detail.

Notably, results obtained at the single-gene level using curated gene families were partially reproduced with the whole-genome approach applied to random sub-cohorts, particularly for genes involved in the TMA-methane metabolic axis and propanol production (Supplementary Fig. 4). This partial overlap supports the accuracy of the single-gene analysis. However, the lack of complete concordance may be attributed either to the reduced sample size in the co-assemblies or to mismatches between our custom gene families and the broader Kegg Orthology (KO) annotations required to detect the same functional signals.

Importantly, we found that metabolic genes associated with MASLD exhibit a highly accessory profile within the GM (Fig. 5). Butyrate-producing genes are highly accessory in nearly half of the most abundant GM genera, while SCA-producing genes display an even more pronounced accessory pattern (Fig. 5A). Among the two main butyrate biosynthesis routes, genes from the butyryl-CoA pathway are more pervasive in the GM than those from the butyryl-P pathway. Notably, *Phocaeicola* and *Bacteroides* - the two most abundant GM genera-were excluded from this analysis due to a limited number of high-quality genomes passing our completeness filters, despite their extensive genomic representation in the UHGG. Interestingly, the two key genes in the butyryl-CoA pathway (*bcd* and *but*) are core in *A. rectalis*. Nonetheless, some strains of this known butyrate-producing species also harbor genes for ethanol and propanol production, which our findings suggest may contribute to MASLD onset. This highlights a critical point: taxonomic signatures alone are insufficient for associating bacterial clades with disease. Genome plasticity introduces substantial strain-level metabolic variability in the GM, leading to distinct phenotypic traits even within the same species: an effect well-documented in both pathogenic and commensal *E. coli* strains^93,94^.

Further supporting their mobility, candidate genes are frequently encoded in plasmids (Fig. 5C). Among these, ethanol formation genes (*aldhB*, *yahK*, and *ahr*) are the most widely distributed, while butyrate- and propanol-related genes are found in plasmids less frequently. Since non-essential, adaptive metabolic pathways are often encoded in plasmids across many bacterial species^94–96^, their presence likely drives phenotypic differences in MASLD without altering the overall species composition detected by conventional metataxonomic marker genes. If MASLD is actually driven by bacterial metabolism, strains from different species, but harboring similar metabolic pathways, may produce similar liver-impacting metabolites, potentially contributing to MASLD pathogenesis regardless of their specific taxonomic position.

## 4. Conclusions

Our study reveals that MASLD is marked by distinct geno-metabolic alterations in the GM that are not captured by conventional taxonomic profiling. Specifically, we observed an increase in genes involved in the production of SCAs and TMA, coupled with a depletion of genes responsible for butyrate and methane production. These results confirm multiple previous findings and suggest a broader metabolic shift that may contribute to the pathogenesis of MASLD, with the displacement of beneficial butyrate-producing taxa such as *Agathobacter rectalis*. Importantly, our work underscores the value of analyzing metagenomes at gene level to extract additional phenotypic signatures beyond what 16S rRNA-based or marker gene-based approaches can offer. By integrating metabolic pathway analysis with the isolation of specific gene families, and by examining gene abundance, accessory gene profiles and plasmid-mediated gene mobilization, we have significantly advanced our understanding of bacterial metabolism in MASLD. This comprehensive approach not only enhances our understanding of the microbial etiology of MASLD, but also paves the way for improved bacterial analysis in clinical settings, offering new avenues for diagnosis and therapeutic intervention in liver pathologies.

## Materials and Methods

### A. Downloading of sequencing metagenomic data

A total of 554 metagenomic sequencing libraries generated from the fecal samples of MASLD and healthy subjects were downloaded from the SRA with SRA Toolkit’s *fastq-dump*^97^. These samples were retrieved using the accession numbers corresponding to five different studies, and they were stratified in three MASLD cohorts, as detailed in Table 1. Information on the collection and processing of fecal samples, bacterial DNA extraction and sequencing protocols can be found in the original studies.

**Table 1:**
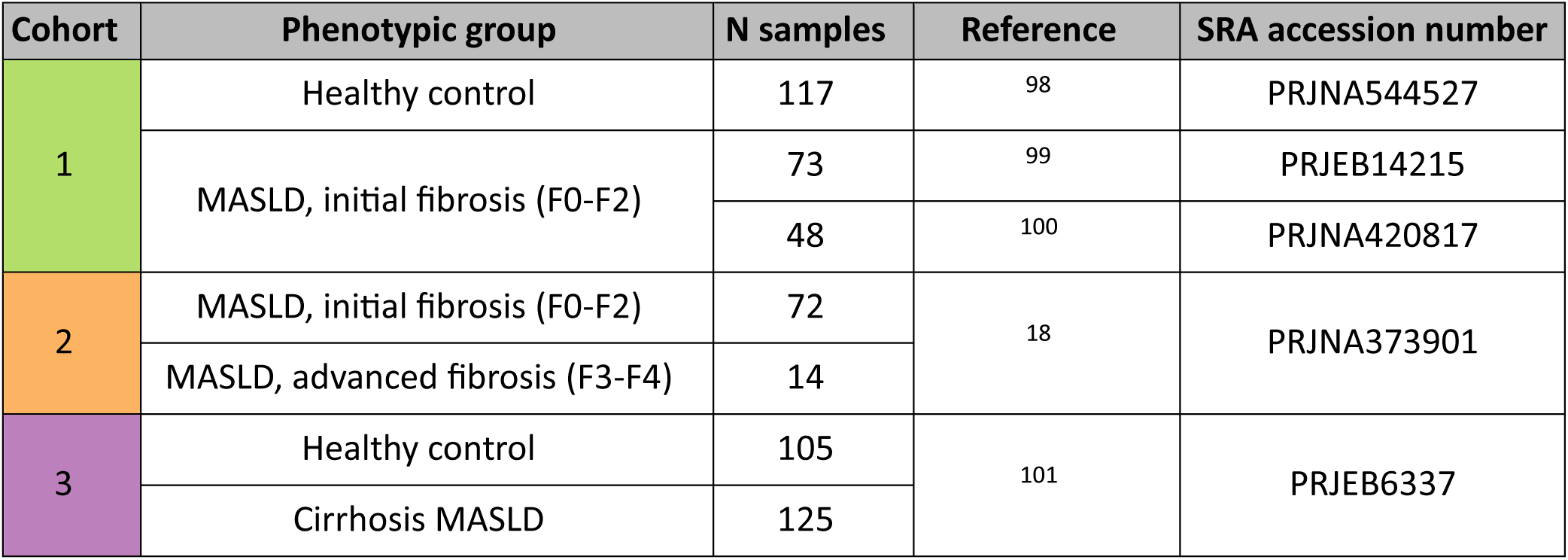
MASLD patient cohorts analyzed in this study. Each cohort (column 1) was divided into two phenotypic comparison groups: control vs. disease (column 2). The cohorts include *N* MASLD patients and corresponding whole-metagenomic stool samples derived from them (column 3). The disease group in Cohort 1 consists of 73^99^ and 48^100^ MASLD patients with initial fibrosis. Cohort 2 was divided into initial fibrosis (control) and advanced fibrosis groups. Patients were originally recruited in the listed studies (column 4), with metagenomic data and medical metadata downloaded from the indicated SRA accession numbers (column 5).

| Cohort | Phenotypic group | N samples | Reference | SRA accession number |
| --- | --- | --- | --- | --- |
| 1 | Healthy control | 117 | 98 | PRJNA544527 |
|  | MASLD, initial fibrosis (F0-F2) | 73 | 99 | PRJEB14215 |
|  |  | 48 | 100 | PRJNA420817 |
| 2 | MASLD, initial fibrosis (F0-F2) | 72 | 18 | PRJNA373901 |
|  | MASLD, advanced fibrosis (F3-F4) | 14 |  |  |
| 3 | Healthy control | 105 | 101 | PRJEB6337 |
|  | Cirrhosis MASLD | 125 |  |  |

Additionally, we retrieved the medical metadata associated with each sample donor to confirm their MASLD status and classify samples into two phenotypic groups of comparison within each cohort. Samples lacking a clearly defined MASLD status were excluded from this study. All data was downloaded between March 2022 and March 2024

### B. Description of MASLD patient cohorts

Cohort 1 consists of two sets of European, obese MASLD patients aged 20-64 years with initial fibrosis (F0-F2) and varying degrees of steatosis: 73 patients from Hoyles *et al.*^99^ and 10 patients from Mardinoglu *et al.*^100^. In the latter group, 48 fecal samples were collected at five time points during the study (two samples from day 1 were not obtained). Diagnoses were confirmed through hepatic biopsy performed during obesity surgery, and none of the subjects had alcoholic liver disease, hepatitis or diabetes. The control group for comparison includes 117 healthy patients from Poyet *et al.*^98^. Cohort 2 comprises 86 American MASLD patients from Loomba *et al.*^18^, including 72 with initial (F0-F2) and 14 with advanced fibrosis (F3-F4). Diagnoses were confirmed through hepatic biopsy and magnetic resonance imaging, and none of the patients had alcoholic liver disease, hepatitis or diabetes. Note that in this study, the initial fibrosis group is referred to as “healthy” for group comparison purposes, despite including patients with early-stage MASLD. Cohort 3 includes 230 Han Chinese patients from Qin *et al.*^101^: 125 diagnosed with cirrhotic MASLD and 105 with no hepatic pathology. None of these subjects had hepatitis. All cohorts consist of one metagenomic stool sample per patient, except for the 48 samples collected at multiple time points from 10 patients in the study from Mardinoglu *et al.*^100^.

### C. Quality control of metagenomic reads

Sequencing reads were trimmed to remove Illumina adapter remnants and low-quality regions (<Q25) using BBDuk from the BBTools suite (version 37.62)^102^. Read quality was assessed with FastQC (version 0.12.1)^103^ and summarized using MultiQC (version 1.22.3)^104^.

### D. Taxonomic signatures associated with MASLD

To determine whether any bacterial clade is systematically increased or decreased in MASLD across the three cohorts, we profiled the taxonomic composition of GM communities in fecal samples using MetaPhlAn (version 4.0.6), which uses a wide-range of clade-specific maker genes to estimate taxonomic abundances^24^. The diversity of these markers allows MetaPhlAn to provide highly accurate taxonomic classification across a wide spectrum of bacterial clades, from the species level to broader taxonomic ranks, without being limited to 16S rRNA alone.

For this study, we used the database version *mpa_vOct22_CHOCOPhlAnSGB_202212*, downloaded on August 22, 2023, and added the *--ignore_eukaryotes* flag to skip profiling eukaryotic organisms. Post-processing of the MetaPhlAn output involved a) removing low-abundant clades (i.e., those with an average relative abundance of 0 in at least one cohort), b) excluding taxa predicted as “unclassified”, and c) verifying that excluded clades were not solely present in only one of the comparison groups, in order to avoid removing what could be very clear taxonomic markers. To identify the most abundant clades present in the GM, we selected genera with an average relative abundance above 0.5% (Fig. 1C) and species above 0.2% (Fig. 1E). Notedly, in this step we faced an issue regarding the status of taxonomic assignments, as several species required manual updates to their nomenclature assigned by MetaPhlAn. Specifically, *Eubacterium rectale* was reassigned to *Agathobacter rectalis*^88^, *Ruminococcus torques* to *Mediterraneibacter torques*^105^ and *Roseburia faecis* to *Agathobacter faecis*^106^.

### E. Isolation of target gene families in the GM

We constructed families of the human GM metabolic genes encoding the enzymes involved in the production of butyrate, ethanol, propanol, TMA, methane and their precursor metabolites, as represented in Figs. 2A, 3A and 4A. The Unified Human Gastrointestinal Genome (UHGG, version 2.0.2)^81^ served as the reference database for this purpose. This resource, comprising 289,232 bacterial and archaeal genomes and more than 665 million proteins, represents the most comprehensive collection of characterized human GM genes to date. Each gene family was built by retrieving a seed set of sequences based on gene and protein annotation from the UHGG, assessing their similarity with lab-validated enzymatic sequences when available. Validated gene sequences were retrieved from MetaCyc^107^ and used as controls to assess their phylogenetic proximity to the retrieved gene seeds.

Gene families were aligned using MAFFT (version 7.271)^108^ with options *“--localpair”* and *“--maxiterate 1000”* and phylogenetic trees were constructed with IQ-TREE (version 2.0.3)^109^ with the ultrafast bootstrap option (1000 bootstraps)^110^ to assess branch support. The best fitting model for each metabolic gene was estimated using ModelFinder Plus^111^, according to the Bayesian information criterion. Low phylogenetic distances between control and retrieved sequences served as a criterion for defining gene families. Duplicated sequences, gene fragments and phylogenetically distant genes were excluded after inspecting the trees. To improve gene identification and classification, we then built profile Hidden Markov Model (HMM) for each target gene family using *HMMER hmpress* (version 3.4)^112^. MetaCyc^107^ and KEGG^113^ were also used to validate gene and enzyme names, KO groups, the biochemical reactions involved in the formation of each metabolite, and their enzyme commission (EC) numbers. The same protocol was applied to isolate five USCGs used as controls. These USCGs and their encoded proteins were: *argS* (arginyl-tRNA synthetase), *dnaA* (chromosomal replication initiator protein DnaA), *rpoA*, *rpoB* and *rpoC* (alpha, beta and gamma subunits of the DNA-directed RNA polymerase, respectively).

### F. Analysis of gene abundance by read-based metagenomic quantification

We quantified the abundance of target genes in the metagenomic samples from the three cohorts by aligning the quality-controlled reads against the isolated gene families using DIAMOND (version 2.0.14)^114^. Alignments were filtered to retain only the best match for each read and gene family sequence by using the *“--max-hsps 1”* option, with an E-value threshold of <1E-10. These parameters were based on experimentally validated recommendations for detecting genes in fecal metagenomic samples with DIAMOND^115^. The total number of aligned reads for each gene family was normalized using a variation of the RPKM formula, referred to as RPKSM (reads per kilobase per size per million reads). This measure was used as an indicator of gene abundance (Figs. 2B, 3B-C and 4B-D; and Supplementary Figs. 2 and 3). It is defined as the number of aligned reads per kilobase per gene family size per million reads, to account for variations in gene length, gene family size (i.e., the number of sequences within each gene family) and library size, respectively.

### G. Whole metagenomic analysis

We applied a whole-metagenome analysis workflow to validate our results at single-gene level. To this end, SqueezeMeta (version 1.6.0)^116^ was used to predict both the taxonomic and functional composition of the metagenomic samples from the three cohorts. Due to computational limitations, it was not feasible to co-assemble all available samples, so approximately 24-30 samples per cohort were randomly selected, with equal representation from each phenotypic group. Briefly, SqueezeMeta assembled the metagenomes from each sub-cohort into a single co-assembly using MEGAHIT (version 1.2.9)^117^ by pooling the sequencing reads from every sample. ORFs were then predicted from contigs^118^, and reads from individual samples were mapped back against the co-assembly to quantify the abundance of each KO group per metagenome. We further processed these results with SQMtools (version 1.6.0)^119^ to identify the most differentially abundant KOs between phenotypic groups within each cohort through DESeq2 (version 1.34.0)^120^, applying a log fold-change >2 and a p-adjusted value <0.05 as thresholds (Supplementary Fig. 4).

### H. Presence of candidate metabolic genes across the human GM genomes and annotated plasmids

To determine whether candidate metabolic genes involved in MASLD were core or accessory across the human GM, their presence was quantified in both UHGG genomes and curated RefSeq200 plasmids^121^. Curation involved eliminating sequences corresponding to partial plasmid DNA sequences, unassignable hosts, or PacBio internal control sequences, as done previously^16^. Since many UHGG genomes originate from metagenome-assembled or single-cell-amplified genomes, they may be incomplete, potentially leading to false positive annotations. To minimize this risk, we included only UHGG genomes with >95% completeness and containing the five USCGs described above. ORFs were predicted using Prodigal (version 2.6.3)^122^, retaining only those covering >90% of the HMM size. We then used HMMER *hmmscan* with the following parameters: protein-pHMM coverage threshold >90%, E-value <0.01 and independent E-value (i-E value) <0.01, strictly enforcing the 90% threshold to avoid capturing ORFs with similar protein domains. This approach yielded 31,227 genomes classified as complete and containing the five USCGs.

To classify genes as core or accessory, we defined their presence as the percentage of strains within a given genus and species that encode the gene. Genes were categorized as core if present in >80% of congeneric (Fig. 5A) or conspecific (Fig. 5B) genomes, as accessory if present in 20-80%, and as highly accessory if present in <20% genomes. Only the most abundant clades in the GM with >100 genomes meeting the completeness criteria were included. Additionally, we assessed whether these genes could be encoded in annotated plasmids. Using the same constraints, we predicted ORFs from the RefSeq200 plasmid collection, which includes 23,309 plasmids encoding over 2 million ORFs. ORFS were retained if they met the >90% HMM size threshold, with *hmmscan* parameters identical to those applied in the UHGG genome analysis (Fig. 5C).

### I. R packages and statistical analysis

Data manipulation was performed using R (version 4.1.3) and the *tidyverse* package (version 1.3.2)^123^. Figures were generated with the R packages *ggplot2* (version 3.4.2)^124^, *ggpubr* (version 0.6.0)^125^ and *pheatmap* (version 1.0.12)^126^. Mann-Whitney tests with a Benjamini-Hochberg adjustment for multiple testing were applied on the relative abundance of bacterial clades (Figs. 1D and 1E) and gene abundance (Figs. 2B, 3B-C, 4B-D, and Supplementary Figs. 2 and 3) between phenotypic groups in the metagenomic fecal samples of each cohort to assess the significance of the differences. Statistical analysis and graphical representation on the plots were conducted using the R package *rstatix* (version 0.7.2)^127^.

