## Supplementary Material for "Bacterial metabolic signatures in MASLD predicted through gene-centric studies in stool metagenomes"

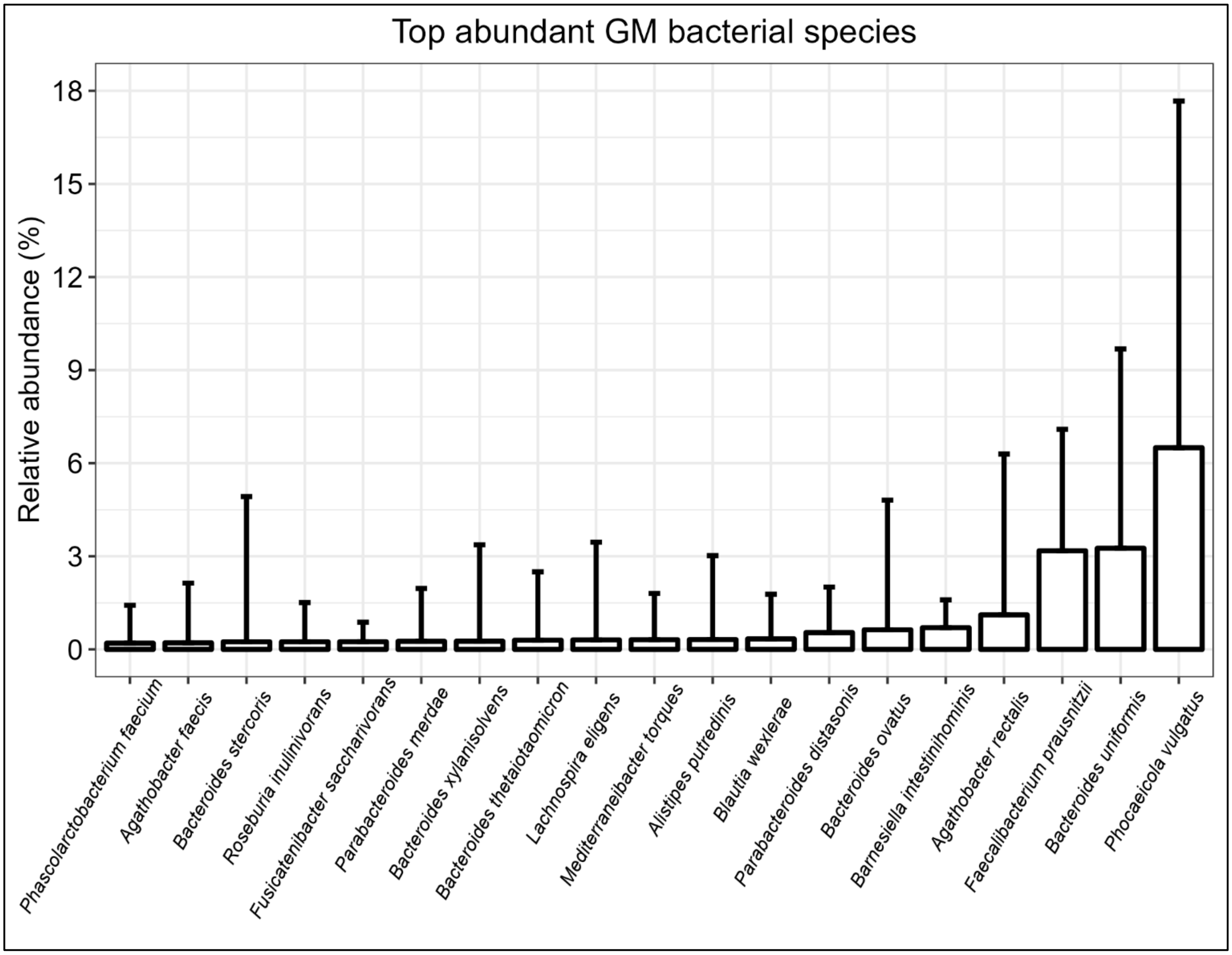


**Supplementary Figure 1. Top abundant GM species** (average absolute abundance >0.2%)

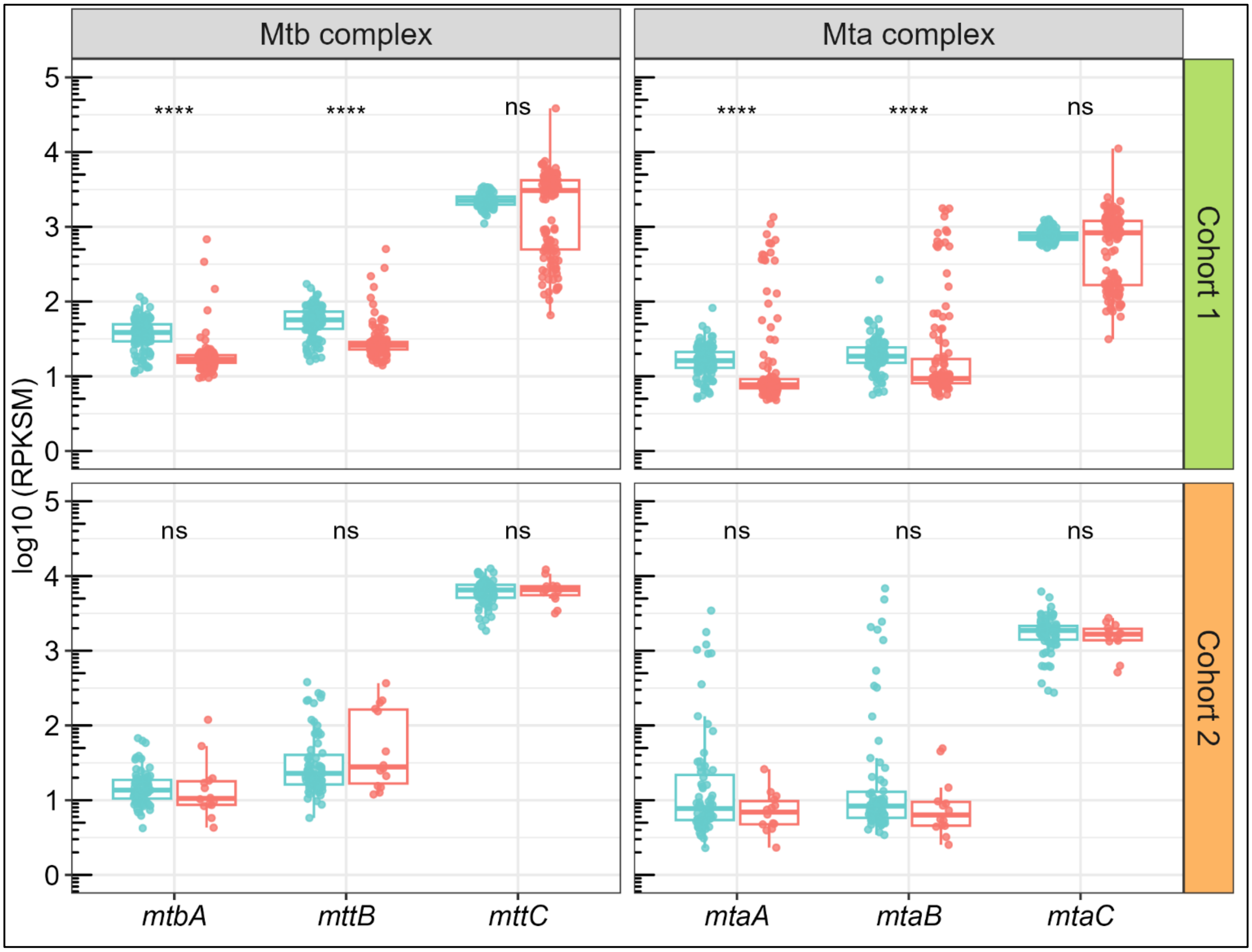


**Supplementary Figure 2. Abundance of genes coding for the Mtb and Mta complexes in patient cohorts.** Boxplot organization and statistical tests were conducted as in Fig. 2B.

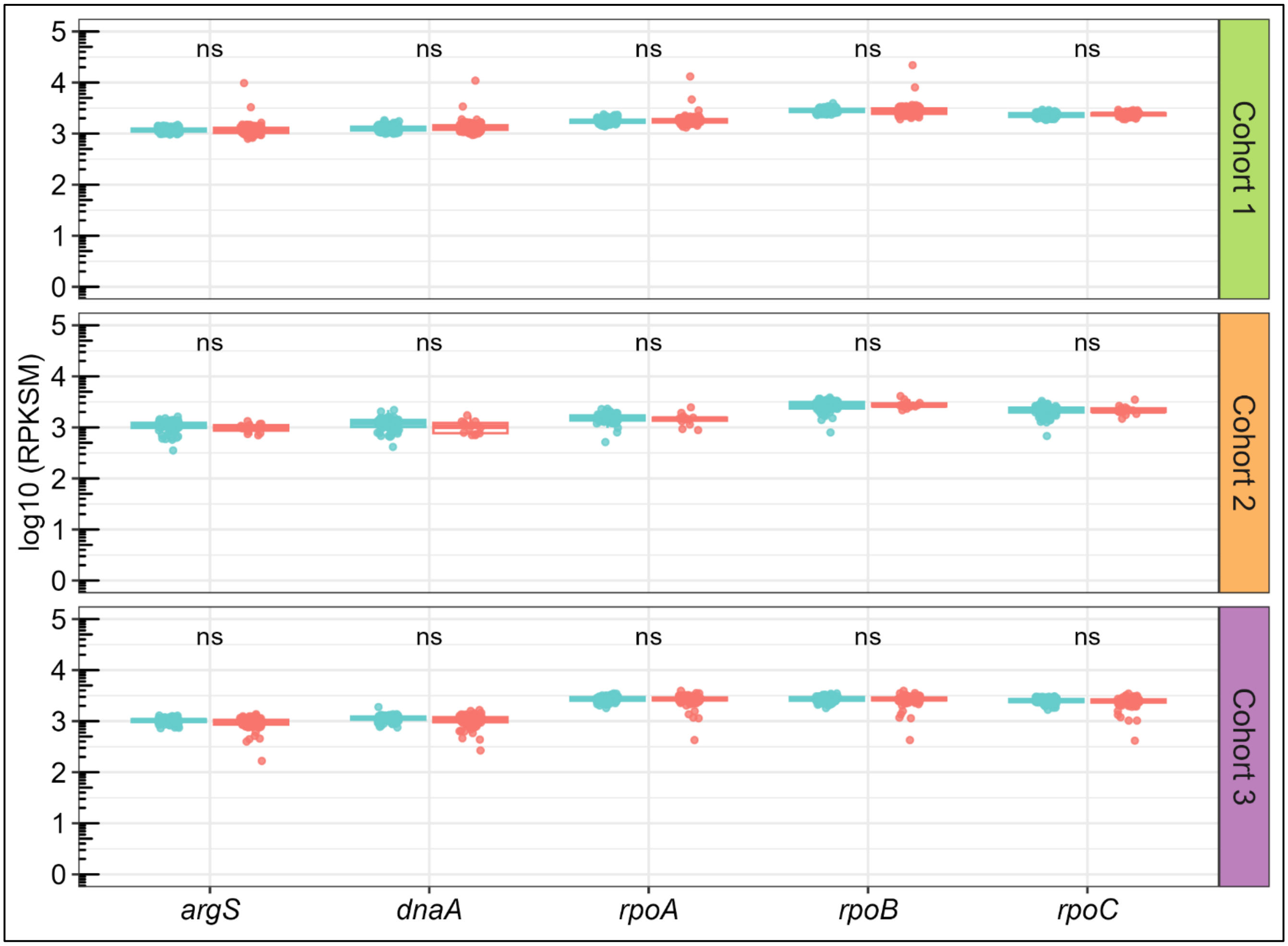


**Supplementary Figure 3. USCGs in patient cohorts.** Abundance of universal, single-copy marker genes in the three cohorts. Boxplot organization and statistical tests were conducted as in Fig. 2B. *argS* encodes for the arginyl-tRNA synthetase*, dnaA* encodes for the chromosomal replication initiator protein DnaA and *rpoA/B/C* encode, respectively, the alpha, beta and gamma subunits of the DNA-directed RNA polymerase.

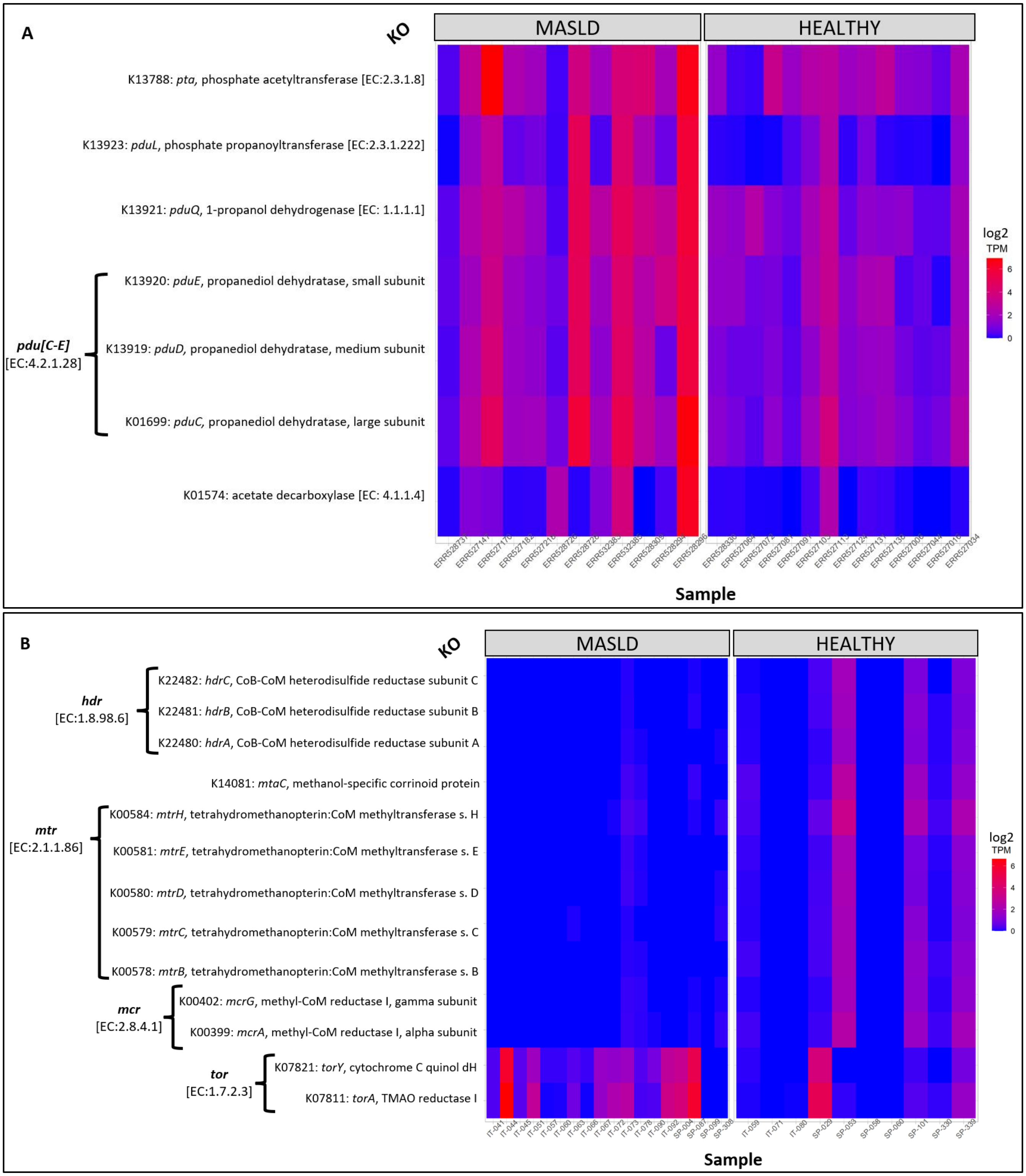


**Supplementary Figure 4. Abundance of KOs predicted from the co-assembled sub-cohorts with SqueezeMeta.** Heatmaps represent KOs involved in the **(A)** propanol-formation and **(B)** TMA-methane metabolic pathway in sub-cohorts 3 and 1, respectively. Genic abundance is expressed in tags per million (TPM), in log2 scale. x-axis indicates samples from both comparison groups and y-axis indicates KO groups, EC numbers, and associated gene and enzyme names according to KEGG. Only KOs with log fold-change >2 and a p-adjusted <0.05 are represented.
